# TNBC response to paclitaxel phenocopies interferon response which reveals cell cycle-associated resistance mechanisms

**DOI:** 10.1101/2024.06.04.596911

**Authors:** Nicholas L Calistri, Tiera A. Liby, Zhi Hu, Hongmei Zhang, Mark Dane, Sean M. Gross, Laura M. Heiser

## Abstract

Paclitaxel is a standard of care neoadjuvant therapy for patients with triple negative breast cancer (TNBC); however, it shows limited benefit for locally advanced or metastatic disease. Here we used a coordinated experimental-computational approach to explore the influence of paclitaxel on the cellular and molecular responses of TNBC cells. We found that escalating doses of paclitaxel resulted in multinucleation, promotion of senescence, and initiation of DNA damage induced apoptosis. Single-cell RNA sequencing (scRNA-seq) of TNBC cells after paclitaxel treatment revealed upregulation of innate immune programs canonically associated with interferon response and downregulation of cell cycle progression programs. Systematic exploration of transcriptional responses to paclitaxel and cancer-associated microenvironmental factors revealed common gene programs induced by paclitaxel, IFNB, and IFNG. Transcription factor (TF) enrichment analysis identified 13 TFs that were both enriched based on activity of downstream targets and also significantly upregulated after paclitaxel treatment. Functional assessment with siRNA knockdown confirmed that the TFs FOSL1, NFE2L2 and ELF3 mediate cellular proliferation and also regulate nuclear structure. We further explored the influence of these TFs on paclitaxel-induced cell cycle behavior via live cell imaging, which revealed altered progression rates through G1, S/G2 and M phases. We found that ELF3 knockdown synergized with paclitaxel treatment to lock cells in a G1 state and prevent cell cycle progression. Analysis of publicly available breast cancer patient data showed that high ELF3 expression was associated with poor prognosis and enrichment programs associated with cell cycle progression. Together these analyses disentangle the diverse aspects of paclitaxel response and identify ELF3 upregulation as a putative biomarker of paclitaxel resistance in TNBC.

## INTRODUCTION

Triple negative breast cancer (TNBC) is an aggressive form of breast cancer that affects 10-20% of all breast cancer patients and is characterized by its lack of expression of estrogen, progesterone and HER2 receptors[1]. The standard of care for TNBC patients primarily relies on conventional anthracycline and taxane-based chemotherapy regimens, and few next-generation therapies have shown efficacy in patients with this disease[2]. Paclitaxel, a taxane-based chemotherapeutic commonly used in TNBC treatment[3], targets microtubules to disrupt the formation of the mitotic spindle, resulting in cell cycle arrest and apoptosis. While 22% of TNBC patients treated with paclitaxel achieve pathological complete response, the outcome for those with residual disease is relatively poor[4, 5]. Moreover, paclitaxel monotherapy only achieves a median 5.5 month progression free survival in patients with locally advanced or metastatic disease[6]. Therefore, there is a need to better understand the molecular basis of paclitaxel response and mechanisms of resistance that may be targeted for therapeutic benefit.

Phenotypic plasticity enables malignant cells to rapidly adapt to therapeutic challenge[7] and can also drive acquired drug resistance[8]. Adaptive responses often involve activation of new transcription factors which in turn upregulate programs that repress immune activation[9], grant tolerance to DNA replication stress[10], or enable evasion of apoptosis [11]. Single-cell RNA sequencing (scRNA-seq) is a powerful approach to investigate the subtle but critical differences in transcriptional landscape that distinguish cellular phenotypic states and to identify molecular programs associated with different therapeutic sensitivities[12]. Single cell methods such as scRNA-seq enable heterogeneous populations to be deconvolved into discrete states to identify the gene regulatory mechanisms that contribute to drug resistance [13, 14].

To elucidate the adaptive responses of TNBC cells to paclitaxel, we performed deep single-cell RNA sequencing of HCC1143 TNBC cells before and after paclitaxel treatment. Paclitaxel induced a range of phenotypic changes, including altered cell cycle phase distribution, increased proportion of multinucleated cells, increased expression of senescence and DNA damage associated biomarkers, and upregulation of interferon-related gene programs. Comparison of gene expression profiles from paclitaxel treated versus IFNB or IFNG treated cells enabled identification of genes that were uniquely upregulated after paclitaxel treatment, including a suite of transcription factors. Functional assessment with siRNA knockdown confirmed that many of these TFs are critical for mediating resistance to paclitaxel. Using live-cell imaging, we probed the temporal dynamics of these functional responses, which demonstrated that knockdown of ELF3, FOSL1 and IRF9 synergize with paclitaxel to slow cell cycle progression. Together, these analyses identify upregulation of ELF3, FOSL1 and IRF9 as important regulators of cell cycle progression that mediate response to paclitaxel, and which may serve as biomarkers of response.

## RESULTS

### Paclitaxel modulates multiple cancer-associated phenotypes

We identified phenotypic changes induced by paclitaxel by treating HCC1143 TNBC cells for 72 hours with paclitaxel, followed by fixation and staining with DAPI (DNA), CellMask (cytoplasmic marker), Tubulin Beta 3 (TUBB3, microtubule component), p16/p15 (senescence) and cPARP (DNA damage induced apoptosis) (**Figure 1A**). We quantified total DAPI intensity to assess cell cycle status[15] and observed two distinct peaks in the DMSO treated sample, representing diploid (G0/G1, mode = 119 A.U) and tetraploid (G2/early M, mode = 222 A.U.) states associated with cycling cells (**Figure 1B**). Intermediate paclitaxel doses (0.01nM-1nM) resulted in an enrichment of cells in the diploid to sub-diploid range, consistent with paclitaxel’s known side-effect of chromosomal disruption[16, 17]. The highest paclitaxel dose tested (81nM) resulted in an increased fraction of cells in diploid and tetraploid states and a broader distribution of nuclear intensities, indicating significant dysregulation of nuclear content. This dysregulation of nuclear content also correlated with a dose-dependent reduction in cell numbers and an increase in the proportion of multinucleated cells (**Figure 1C**). The fraction of multinucleated cells plateaued around 9nM paclitaxel, with ∼25% of surviving cells harboring two or more nuclear structures for all higher dosages.

**Figure 1:**
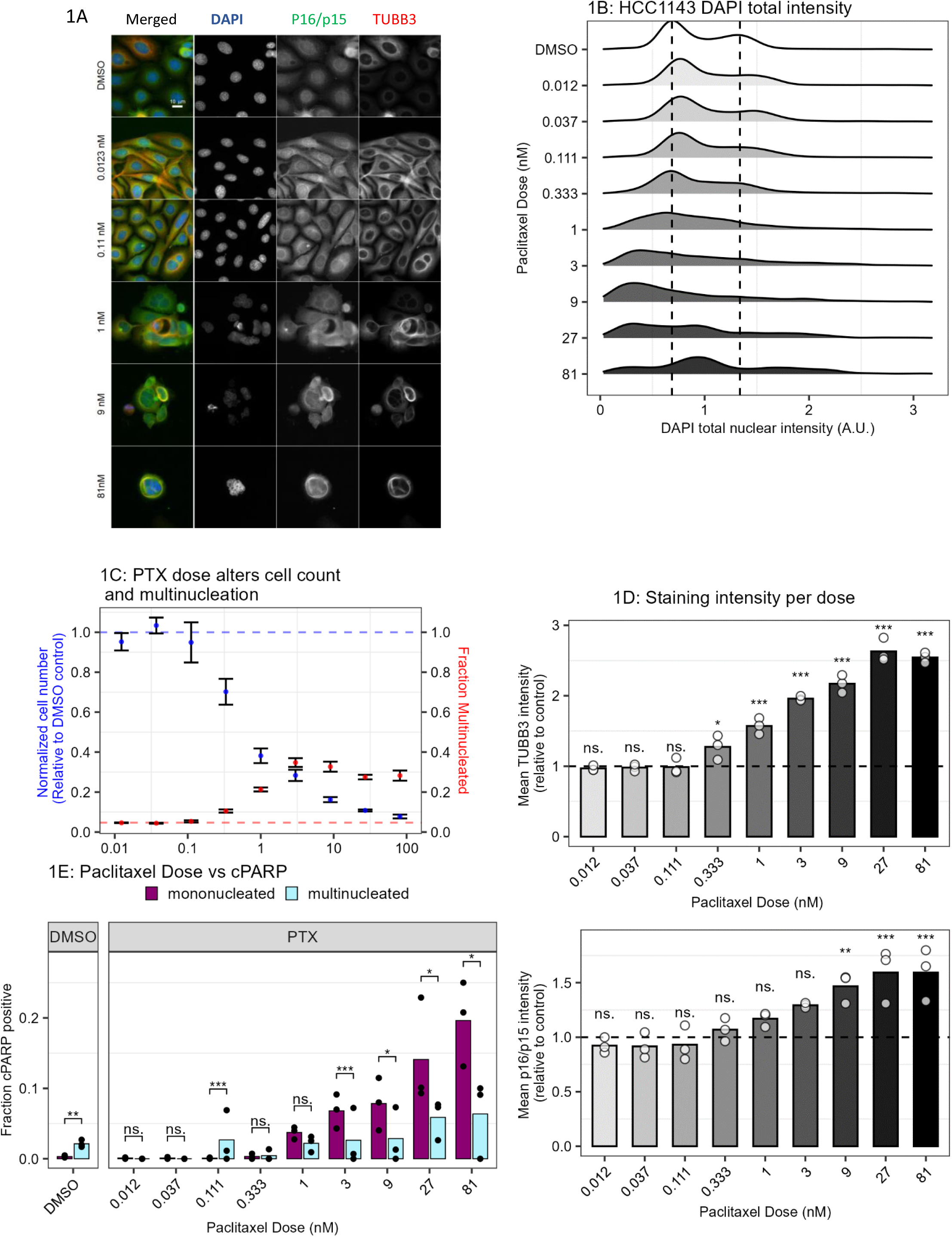
Paclitaxel modulates multiple cancer associated phenotypes. **1A)** Representative fluorescent images showing HCC1143 cells treated with DMSO or Paclitaxel at the listed doses for 72 hours and stained with DAPI, p16-INK4A, and TUBB3. **1B)** Ridgeplot showing impact of paclitaxel treatment on DAPI total nuclear intensity as a proxy for nuclear content. Dashed lines indicate local maxima in the DMSO control condition corresponding with 2N and 4N nuclear state. **1C)** Normalized cell count and fraction of multinucleated cells for HCC1143 treated with serial titration of Paclitaxel for 72 hours. Error bar indicates SEM across 6 replicates. **1D)** Barplots showing mean TUBB3 and p16/p15 cytoplasmic staining intensity for triplicate wells of HCC1143 treated with a range of Paclitaxel and normalized to paired DMSO control (horizontal line). Significance assessed with Dunnett’s test. **1E)** Barplot comparing the fraction of cPARP positive cells for mononucleated (magenta) versus multinucleated (cyan) cells within the same treatment condition. cPARP positive threshold was set to the 99th quantile of DMSO treated cells total cPARP nuclear intensity (Supplemental Figure 1C). Significance assessed with proportions test. For all statistics: * = p<0.05, ** = p<0.01, *** = p<0.001.

We further assessed adaptive cellular responses by analyzing biomarkers associated with senescence (p16/p15), DNA damage induced apoptosis (cPARP), and microtubule component (TUBB3). TUBB3 overexpression has been associated with resistance to multiple microtubule targeting drugs, and consistent with this, we found a dose-dependent relationship between TUBB3 expression and paclitaxel concentration[18, 19]. There was also a positive association between both cytoplasmic and nuclear p16/p15 staining and paclitaxel dose (**Figure 1D**). Additionally, we observed a strong correlation between p16/p15 and TUBB3 expression at the single cell level across paclitaxel concentrations, suggesting that the TUBB3 highly expressing cells represent a senescent subpopulation of cells (**Supplemental Figures 1A,1B**, Pearson correlation = 0.70, r^2 = 0.48). Increasing doses of paclitaxel induced a corresponding increase in the fraction of cPARP positive cells (DMSO: 6%, 81nM Paclitaxel: 28% cPARP positive), indicating induction of DNA damage driven apoptosis (**Figure 1E, Supplemental Figure 1C**). Higher paclitaxel doses resulted in a significantly higher proportion of mononucleated cells staining positive for cPARP as compared to multinucleated (19.4% mononucleated cells and 7.1% multinucleated cells cPARP positive at 81nM paclitaxel, proportions test p = 0.017), suggesting that multinucleated cells are less likely to undergo DNA damage-induced apoptosis. Together this suggests that the multinucleated cells that survive paclitaxel treatment are cell cycle arrested and also less likely to undergo DNA damage-induced apoptosis than mononucleated cells.

### Cells surviving paclitaxel treatment halt cycling and upregulate interferon response genes

To assess paclitaxel-induced molecular programs, we performed 10X Genomics single-cell whole transcriptome sequencing of HCC1143 cells treated with either DMSO vehicle control or 1nM paclitaxel for 24 hours or 72 hours (**Figure 2A**). After quality control filtering that required cells to have a minimum of 3000 unique genes and a maximum of 25% mitochondrial counts, we recovered 3194 total cells (513 – 1106 cells per condition) with a mean UMI count of 63,668 (**Supplemental Figure 2A**).

**Figure 2:**
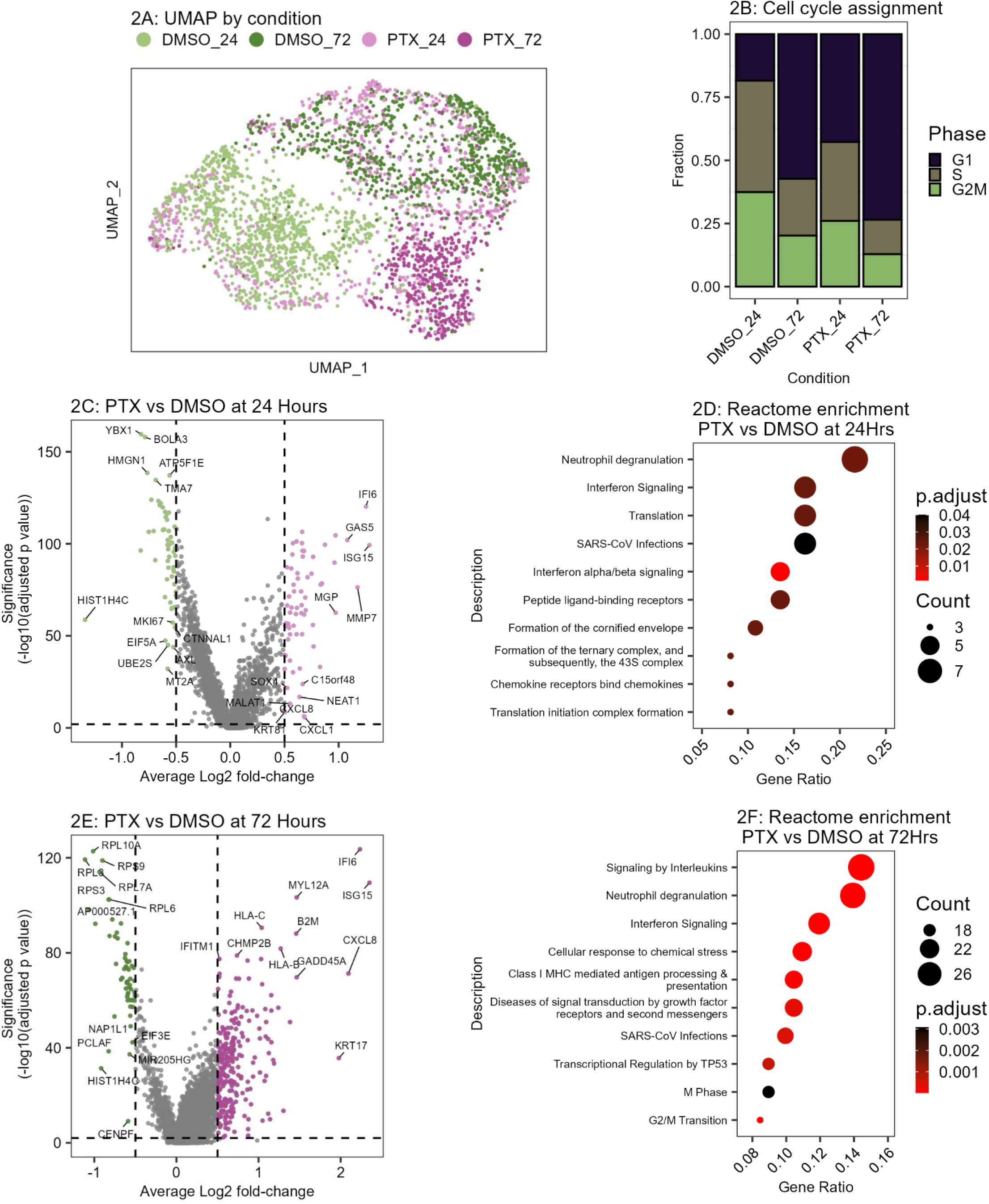
Cells surviving paclitaxel treatment halt cycling and upregulate interferon response genes. **2A)** UMAP color coded by treatment condition. DMSO_24 = 0.1% DMSO for 24 hours, DMSO_72 = 0.1% DMSO for 72 hours, PTX_24 = 1nM Paclitaxel for 24 hours, PTX_72 = 1nM Paclitaxel for 72 hours. **2B)** Barplot showing proportion of each condition assigned to G1, S, or G2M cell cycle state based on transcriptional profile. **2C,2D)** Volcano plot of differentially expressed genes for Paclitaxel treatment versus DMSO at 24 (2C) and 72 (2D) hours. Differentially expressed genes (black) determined with cutoffs of Benjamini Hochberg corrected p<0.05 and absolute Log2FoldChange > 0.5. **2D)** Reactome pathway enrichment results for genes significantly upregulated after paclitaxel treatment at 24 hours. Size indicates the number of genes upregulated within the pathway, color indicates significance. **2E)** Volcano plot of differentially expressed genes for Paclitaxel treatment versus DMSO at 72 hours. Differentially expressed genes (black) determined with cutoffs of Benjamini Hochberg corrected p<0.05 and absolute Log2FoldChange > 0.5. **2F)** Reactome pathway enrichment results for genes significantly upregulated after paclitaxel treatment at 72 hours. Size indicates the number of genes upregulated within the pathway, color indicates significance.

We examined drug-induced changes in cell cycle distribution by assigning cell cycle status to each individual cell using aggregate expression of canonical gene programs for S and G2/M[18,19]. In agreement with our imaging results, we observed an enrichment in the fraction of G1 cells after paclitaxel treatment as compared to time-matched vehicle control (**Figure 2B**). Unsupervised clustering tended to group cells by treatment condition and cell cycle phase (**Supplemental Figures 2B, 2C, 2D**).

We analyzed time-matched conditions to identify significantly differentially expressed genes induced by paclitaxel treatment (Wilcoxon rank sum test, absolute log2 fold-change > 0.5, Benjamini Hochberg FDR < 0.01). This revealed a time-dependent change in molecular programs with 66 significantly upregulated and 57 significantly downregulated genes after 24 hours of paclitaxel treatment, and 256 significantly upregulated genes and 58 significantly downregulated genes after 72 hours (**Supplemental Figure 2E**). Reactome pathway enrichment analysis revealed that the significantly upregulated genes from the 24-hour paclitaxel treated sample were enriched for multiple programs related to Interferon Signaling and Translation (**Figures 2C-D**, **Supplemental Data 2**). Programs uniquely upregulated after 72 hours of paclitaxel treatment include Response to Chemical Stress, Cell Cycle Progression and Antigen Processing-Cross presentation (**Figure 2E-F**). The ontologies enriched after 72-hour paclitaxel treatment had low overlap with those at 24 hours (Jaccard Index = 0.023, **Supplemental Figure 2F**). Notably, the Neutrophil Degranulation pathway was significantly enriched at both time points, with upregulated genes related to antigen presentation (HLA-B, HLA-C, B2M) and differentiation (CD47, CD55, CD59, CD63). Paclitaxel treatment also induced significant upregulation of the pro-tumorigenic chemokines CXCL1 and CXCL8 (**Supplemental Figure 2G**)[20–23]. Together this shows that TNBC cells that survive paclitaxel treatment have altered surface marker expression and produce tumor supportive chemokines.

### Paclitaxel response activates canonical interferon response genes

Despite gene enrichment consistent with interferon response, the paclitaxel treated cells showed no evidence of autocrine signaling, indicating that paclitaxel induces interferon response pathways in a non-canonical manner (**Supplemental Figure 3A**). To disentangle the paclitaxel response signature from true interferon response, we performed a second scRNA-seq experiment with HCC1143 cells that were treated for 72 hours with 7 perturbations that target ligand-receptor pairs known to play an important role in normal and pathological breast tissue[24, 25]: Interferon-Beta (IFNB), Interferon-Gamma (IFNG), Transforming Growth Factor Beta (TGFB), Oncostatin-M (OSM), Lymphotoxin Alpha (LTA), Notch Inhibitor (NOTCHi) and combination of Notch Inhibitor and Interferon-Beta (NOTCHi_IFNB). Cells were treated for 72 hours and then harvested and sequenced with the 10X Genomics scRNA-seq pipeline. After quality control filtering, we recovered 4231 total cells (295 – 725 cells per condition, **Supplemental Figure 3B**).

Overall, the scRNA-seq data revealed that the treated cells largely grouped by perturbation (Normalized Mutual Information = 0.58, **Figure 3A**) and cell cycle state (Normalized Mutual Information = 0.28, **Supplemental Figure 3C**). The IFNB, IFNG, TGFB, NOTCHi and NOTCHi_IFNB conditions all had an increase in proportion of G1 cells compared to control, suggesting these ligands are cytostatic in this cell line (**Supplemental Figure 3D**). Based on the observation that paclitaxel induced Interferon related pathways, we next sought to evaluate the similarity in transcriptional response between paclitaxel and the ligand perturbations. To that end, we computed the differential expression of all genes for each perturbation compared to time-matched vehicle control and then evaluated the pairwise Pearson correlation of log2 fold-change values (**Figure 3B**). The IFNB and IFNG conditions were the most strongly correlated (Pearson correlation = 0.86), indicating a conserved impact on transcription despite acting through different receptors. We found that the 72-hour paclitaxel condition was highly correlated with the interferon treatments (IFNB Pearson correlation = 0.57, IFNG Pearson correlation = 0.47) as compared to the other single-agent perturbations (0.0, 0.11, 0.18, 0.38 Pearson correlation with OSM, LTA, NOTCHi and TGFB respectively).

**Figure 3:**
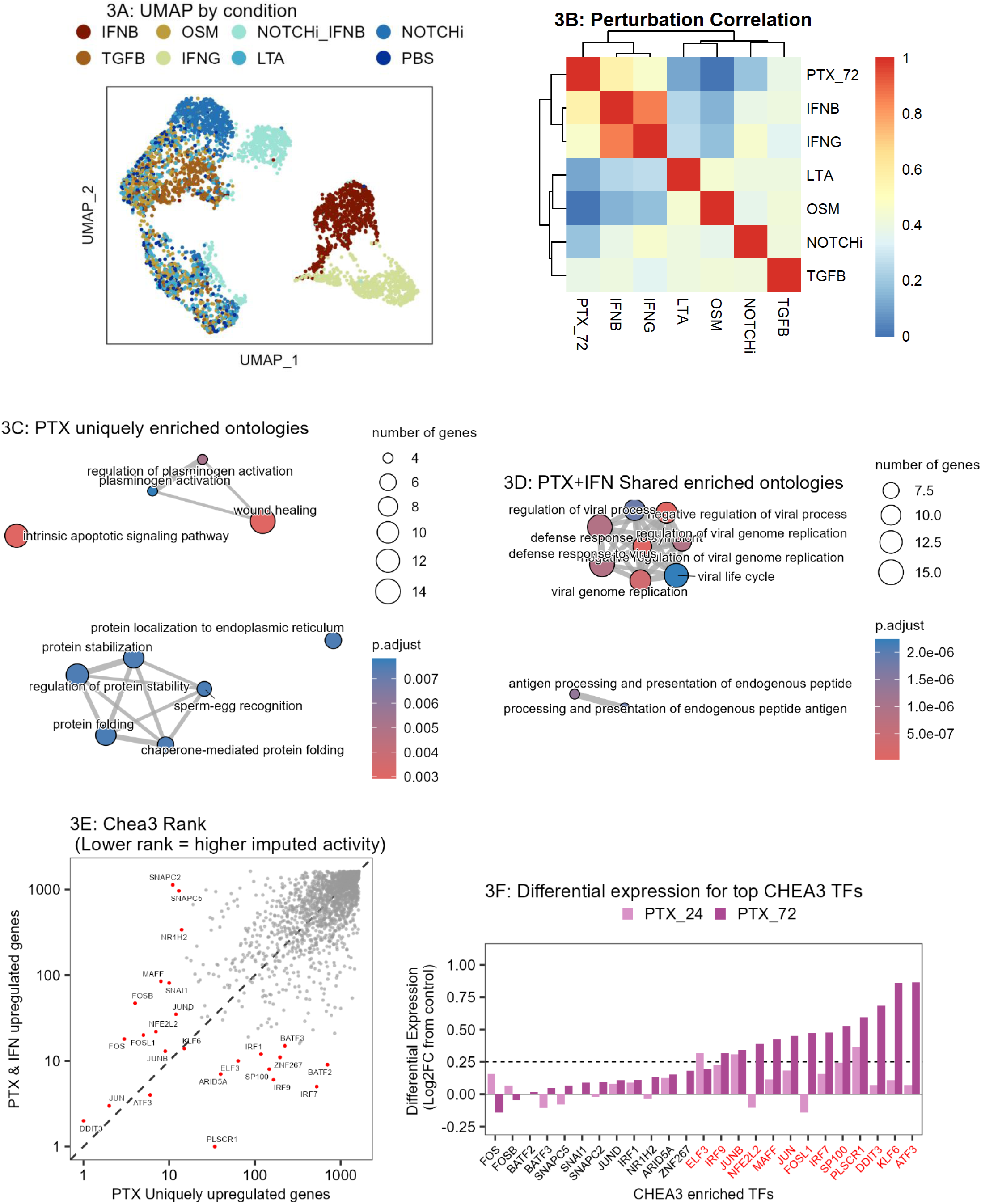
Paclitaxel response activates canonical interferon response genes. **3A)** UMAP showing the scRNA-seq landscape for ligand perturbations. IFNB = Interferon-Beta, OSM = Oncostatin-M, NOTCHi_IFNB = Notch inhibitor + Interferon-Beta, NOTCHi = Notch inhibitor, TGFB = Transforming Growth Factor Beta, IFNG = Interferon-Gamma, LTA = Lymphotoxin-Alpha, PBS = Phosphate Buffered Saline (control). **3B)** Heatmap showing the Pearson correlation for all gene log2 fold-change between perturbation versus time-matched control. Inset number and color indicate correlation. **3C,D)** Gene enrichment map for Paclitaxel uniquely upregulated (3C) and Paclitaxel+Interferon shared upregulated (3D) genes. Color indicates significance, size indicates number of upregulated genes, and lines connect ontologies with shared elements. **3E)** ChEA3 transcription factor enrichment ranks computed from 140 Paclitaxel uniquely upregulated genes (x axis) versus 120 Paclitaxel-Interferon shared upregulated genes (y axis). Lower rank indicates higher imputed activity. TFs to the lower right of the diagonal have higher imputed activity within the PTX+IFN shared upregulated gene set, and TFs to the upper left of the diagonal have higher imputed activity within the PTX uniquely upregulated gene set. **3F)** Bar plot showing Average Log2FC from paclitaxel treated scRNA-seq data for the 24 top ranked transcription factors (intersect of top 15 ranked for PTX unique or PTX shared individually). Transcription factor names in red had differential upregulation (average log2 fold-change > 0.25, FDR < 0.01) at either 24 or 72 hours of paclitaxel treatment compared to vehicle control.

While type 1 and type 2 interferons primarily exhibit antitumor effects through activation of the immune system, some studies have found they have direct effects through induction of cell cycle arrest or apoptosis in malignant cells[26, 27]. To better understand the overlapping transcriptional responses of paclitaxel and interferon, we next sought to differentiate between pathways that were uniquely induced by paclitaxel response or that represent common responses induced by paclitaxel or interferon perturbation. Reactome pathway enrichment analysis revealed that the 140 genes upregulated after paclitaxel treatment but not after IFNG or IFNB (“paclitaxel-unique”) were enriched in molecular programs related to wound healing, protein folding, and intrinsic apoptotic signaling pathway (**Figure 3C**), whereas the 117 genes upregulated by all three treatments (“paclitaxel-shared”) were associated with defense response to virus and antigen presentation (**Figure 3D**).

We hypothesized that the strong overlap in interferon and paclitaxel transcriptional responses was driven by a shared increase in transcription factor (TF) activity through activation of cytosolic DNA sensing pathways[28, 29]. We identified enriched TFs in our paclitaxel-unique and paclitaxel-shared gene signatures using ChEA3[30], which evaluates the expression of gene targets downstream from a TF of interest. DDIT3, JUN, KLF6, and ATF3 emerged as enriched transcription factors across both the paclitaxel-unique and paclitaxel-shared gene signatures (**Figure 3E**). The paclitaxel-unique genes were enriched for TFs in the Immediate-Early Gene family, including JUN (JUN, JUNB, JUND) and FOS (FOS, FOSL1, FOSB)[31]. TFs enriched from the shared gene list were associated with activity of Interferon Regulatory Factors (IRF1/IRF7/IRF9) and Basic Leucine Zipper family (BATF2/BATF3), both related to antiviral response and regulation of antigen-presenting cells[32–34]. The high activity of IRF7 is consistent with activation of the cytosolic nucleotide sensor RIGI, suggesting that the nuclear damage induced by paclitaxel drives an increase in cytosolic RNA or DNA [35].

### Inhibition of paclitaxel-induced transcription factors alters proliferation and nuclear morphology

We used siRNA knockdown in three basal-like TNBC cell lines (HCC1143, HCC1806, MDA-MB-468) to functionally assess prioritized TFs implicated in modulating response to paclitaxel. We nominated a panel of 13 TFs for functional testing, based on ChEA3 analysis and change in gene expression after 24 or 72 hours of paclitaxel treatment (**Figure 3F**, Log2FC > 0.25, Benjamini-Hochberg FDR < 0.01). Most of the TFs included in this panel were either subunits of (ATF3, FOSL1, JUN, JUNB, MAFF)[36] or known interactors with (ELF3, IRF7, DDIT3, NFE2L2)[37–40] the AP-1 transcription factor family. Dysregulation of the AP-1 pathway is associated with multiple tumorigenic phenotypes including enhanced cellular growth, proliferation, and survival[41]. To functionally assess the role of these TFs in paclitaxel response, cells were transfected with siRNA for 24 hours, then treated for 72 hours with paclitaxel or DMSO, and subsequently fixed and stained with DAPI (nuclear marker) and CellMask (cytoplasmic marker). The resultant images were subjected to quantitative image analysis to identify nuclear and cellular masks, followed by quantification of total cell number and fraction of multinucleated cells for each condition.

First, we analyzed the influence of TF knock-down on cell count after 72 hours to evaluate their effects on cell viability. We found that knockdown of 3 of 13 TFs (NFE2L2, IRF7, MAFF) in the absence of paclitaxel significantly reduced cell numbers for at least one cell line (Student’s t-test, p<0.05, **Figure 4A** green bars, **Supplemental Figure 4A**). We then examined the influence of TF knock-down in the presence of paclitaxel to test our hypothesis that upregulation of these TFs mediates adaptive resistance. Knockdown of 5 of 13 TFs (NFE2L2, ELF3, IRF7, FOSL1, PLSCR1) in combination with paclitaxel significantly lowered cell count in at least one cell line compared to paclitaxel alone (Student’s t-test, p < 0.05, **Figure 4A** purple bars, **Supplemental Figure 4A**).

**Figure 4:**
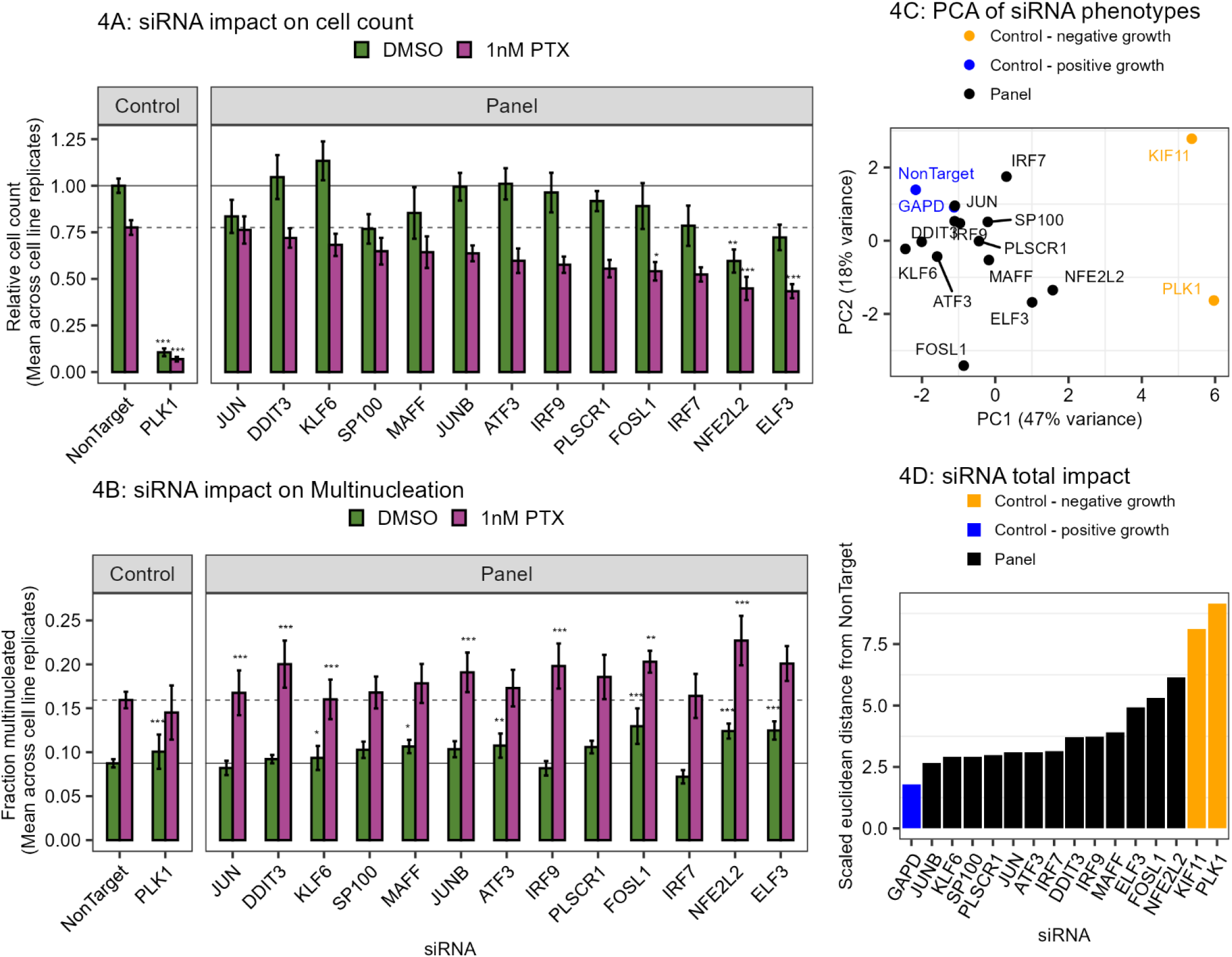
Inhibition of paclitaxel-induced transcription factors alters proliferation and nuclear morphology. **4A-B)** Barplots showing relative cell count (A) and proportion of multinucleated cells (B). Cell count is normalized to the same cell-line DMSO + siNonTarget control. Bars show the mean of three cell lines, and error bar indicates SEM. Relative cell count statistics computed with Fisher’s multi test applied to Two-tailed Student’s T-test per cell line, and fraction multinucleated statistics computed with Fisher’s multi test applied to proportions test per cell line. Heatmap of all values in Supplement 4A, 4B. Not shown: secondary positive growth (siGAPD) and negative growth (siKIF11) controls **4C)** Principal Component results for each siRNA knockdown where each combination of cell line (HCC1143, HCC1806, MDA-MB-468), feature (relative cell count, fraction multinucleated) and condition (DMSO, PTX) is considered a feature (Supplemental Figure 4C). **4D)** The Euclidean feature-distance from NonTarget control for each siRNA. Heatmap of scaled feature values in Supplemental Figure 4C. For all statistics: * = p<0.05, ** = p<0.01, *** = p<0.001.

We hypothesized that these 13 TFs may also be involved in cytokinesis, based on our observation that escalating paclitaxel dose was associated with an increased fraction of multinucleated cells (**Figure 1C**). We found that siRNA knockdown alone caused a significant increase in the fraction of multinucleated cells for 6 of 13 TFs for at least one cell line (NFE2L2, ELF3, SP100, FOSL1, MAFF, and ATF3. Proportions test, p < 0.05. **Figure 4B, Supplemental Figure 4B**). Additionally, knockdown for 11 of 13 TFs in the presence of paclitaxel resulted in significantly increased fraction of multinucleated cells as compared to paclitaxel alone, for at least once cell line. These findings suggest an important role for these transcription factors in maintaining nuclear structure and achieving symmetric cytokinesis in proliferating breast cancer cells.

We comprehensively analyzed the influence of siRNA knockdown across cell lines and drug conditions. Here, we considered each siRNA an independent sample and each combination of cell line (HCC1143, HCC1806, MDA-MB-468), treatment (DMSO, paclitaxel) and phenotype (relative cell count, fraction multinucleated) as 12 independent features (**Supplemental Figure 4C**). Principal component analysis applied to the transformed data revealed that our positive and negative growth controls separated along Component 1 (**Figure 4C**). To identify the TFs that had the greatest overall impact on phenotype, we computed the Euclidean feature-distance (distance for z-scored features) to identify TFs that induced the greatest feature-distance from siNonTarget positive growth control. We found that ELF3, FOSL1, and NFE2L2 knockdown had the largest Euclidean feature-distance (**Figure 4D**) and additionally separated from the rest of the panel via hierarchical clustering (**Supplemental Figure 4C**), indicating that knockdown had a strong effect on both proliferation and regulation of nuclear morphology across the three TNBC cell lines. Protein quantification for ELF3, FOSL1 and NFE2L2 confirmed that the siNonTarget+Paclitaxel induced an accumulation of protein compared to vehicle control, and the targeted siRNA+Paclitaxel reduced protein levels to below the siNonTarget+Vehicle level (**Supplemental Figures 5A-B**).

### ELF3 and FOSL1 mediate cell cycle progression under paclitaxel treatment

Motivated by the observation that many anti-cancer drugs act by targeting the cell cycle, we next explored the influence of prioritized TFs on cell cycle progression by leveraging a genetically engineered HCC1143 cell cycle reporter cell line. The cell cycle state of HDHB-mClover/NLS-mCherry HCC1143 cells can be determined by quantification of relative HDHB-mClover (nuclear translocating cell cycle reporter) intensity within the cytoplasm compared to the nuclear signal marked by NLS-mCherry (stable nuclear localization)[42, 43]. Cells in G1 cell cycle phase have near-equal nuclear and cytoplasmic HDHB-mClover expression, cells in S/G2 cell cycle phases exclude the HDHB-mClover from the nucleus, and cells in M phase concentrate the HDHB-mClover expression to the nucleus. Here we focused on NFE2L2, ELF3 and FOSL1, which induced the largest phenotypic effects; we additionally tested IRF9 which has been previously linked to anti-microtubule chemotherapy resistance [44]. Reporter cells were subjected to siRNA transfection for 24 hours and then treated with either 1nM paclitaxel or DMSO. Treated cells were imaged every 15 minutes for 72 hours. Nuclear and cytoplasmic masks were segmented with custom trained Cellpose[45] models and the resultant data used to classify cells into four ‘phase’ assignments based on HDHB-mClover expression and their number of nuclei (**Figure 5A**). Mononucleated cells were assigned as ‘G1’, ‘S/G2’ or ‘M’ phase and multinucleated cells assigned to either ‘M’ or ‘multinucleated’ phase based on localization of the HDHB-mClover signal (**Supplemental Figure 6A**).

**Figure 5:**
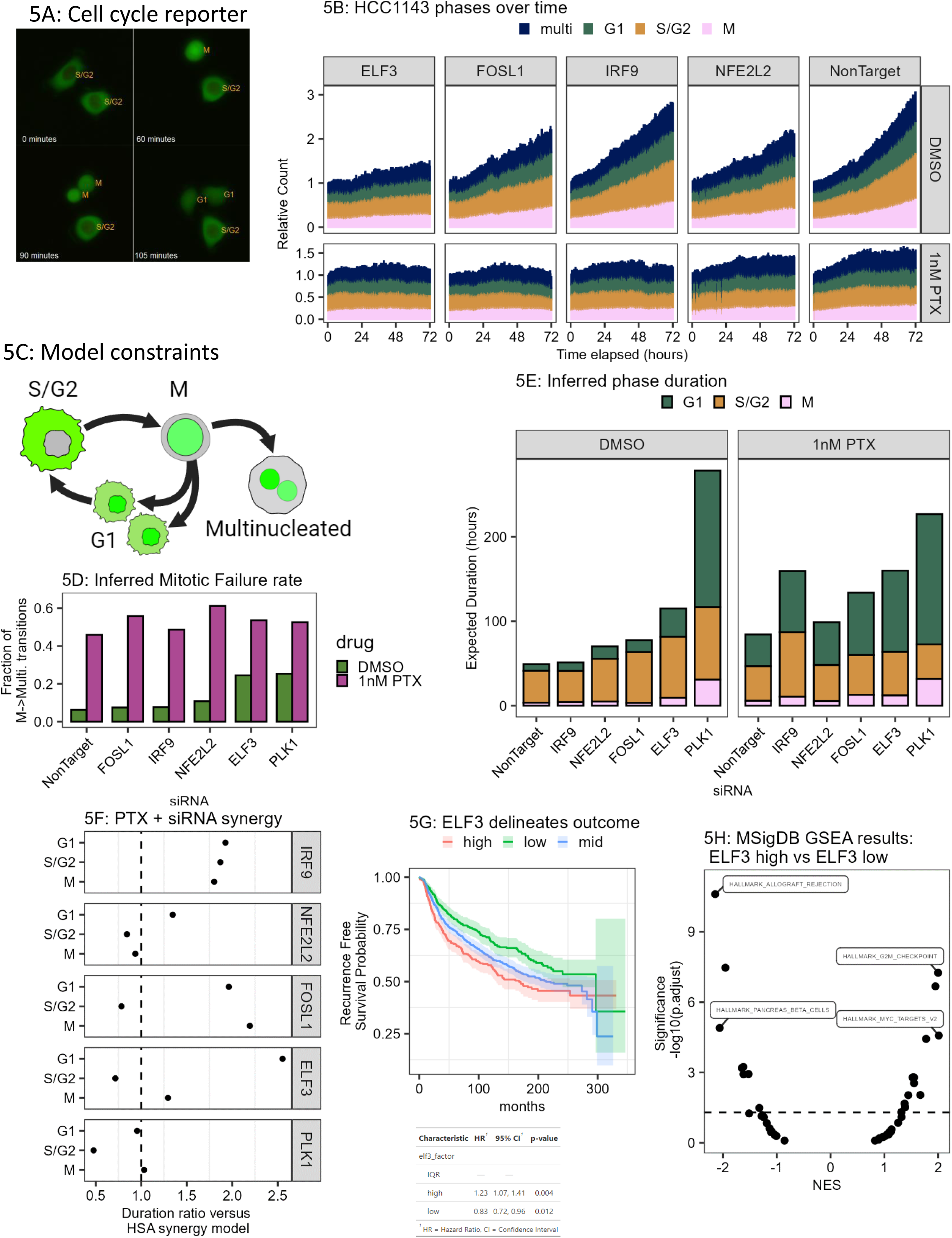
ELF3 and FOSL1 mediate cell cycle progression under paclitaxel treatment. **5A)** Representative images showing the HCC1143 cell cycle reporter line and a mitotic even occurring over 105 minutes. Orange text indicates automatically assigned cell cycle for the processed images. **5B)** Relative (normalized to total cell number at earliest time point) cell count for each phase over time for each siRNA condition +/- paclitaxel (PTX). **5C)** Schematic showing the underlying structure of permitted transitions used in the Markov Model. **5D)** Mitotic failure rate computed from Markov model transition rates. Mitotic failure rate is calculated as the ratio of M->multinucleated transitions divided by the sum of M->G1 and M->multinucleated transition rates. **5F)** PTX + siRNA synergy computed as the ratio of inferred phase duration for combination (siRNA + PTX) versus Highest Single Agent (HSA, highest duration for either siRNA or PTX treatment alone). Value of 1 indicates no change in combination, values greater than 1 indicate synergy and values less than 1 indicate antagonism. **5G)** Overall survival for the Metabric breast cancer cohort striated by ELF3 mRNA expression. High = top quartile of ELF3 expression, IQR = inner quartile range of ELF3 expression, and low = lowest quartile of ELF3 expression. **5H)** MSigDB Gene Set Enrichment (GSEA) results for ELF3 high versus ELF3 group. Horizontal line represents a FDR threshold of 0.05.

As chemotherapeutic drugs often have peak efficacy during a specific cell cycle phase, we next asked whether the combination of paclitaxel treatment and siRNA knockdown altered the dynamics of cell cycle progression. To that end, we trained a Markov Model on the live-cell data, which enabled us to infer transition rates and the average time spent in each of the four phases for a given treatment condition[42, 46]. This approach uses the change in fraction of cells in each cell cycle phase over time (**Figure 5B**) to learn cell cycle-specific transition rates, which represent the fraction of cells that transition from one phase to another phase within a 1-hour timestep (**Supplemental Figure 6B**). We constrained our model such that proliferating cells can either successfully complete the cell cycle or undergo mitotic failure into a permanent multinucleated phase (**Figure 5C**).

We compared the model output to the observed phase counts and found similar trends over time and for all treatment conditions (**Supplemental Figures 6C, 6D**). We used a local polynomial regression (second order LOESS) as a reference and found that our Markov model output compared favorably, with a Root Squared Mean Error (RMSRE) of only 0.0809 in excess of the LOESS fit (**Supplementary Figures 7A, 7B**). Additionally, we used the Chi-squared test to evaluate whether there was a significant compositional difference between the experimental data and Markov output. This approach found that greater than 95% of timepoints had no significant difference for all conditions except for DMSO+siFOSL1 (**Supplemental Figure 7C**). For the DMSO+siFOSL1 condition, the model and experimental data agreed for the first 60 hours but diverged during the last 12 hours, with the model predicting fewer M phase cells than observed in the experimental data.

We next leveraged the model’s learned transition rates to calculate the rate of mitotic failure for each condition and to better understand how the combination of siRNA knockdown and paclitaxel treatment compared to siRNA knockdown alone. The Markov model framework enabled calculation of the mitotic failure rate as the ratio of M->multinucleated transition rate over the sum of M->G1 and M->multinucleated transition rates (**Figure 5D**). siRNA knockdown of ELF3 alone had the largest effect without paclitaxel and increased the mitotic failure rate by 18.1% (DMSO+siNonTarget = 6.4%, DMSO+siELF3 = 24.5%), while inhibition of FOSL1, NFE2L2 or IRF9 had limited effects (DMSO+siFOSL1 = 7.5%, DMSO+siIRF9 = 7.7%, DMSO+siNFE2L2 = 10.8%). Combination siRNA knockdown and paclitaxel treatment resulted in higher mitotic failure rates for ELF3, FOSL1, and NFE2L2 (PTX+siNonTarget = 45.9%, PTX+siELF3 53.6%, PTX+siFOSL1 = 55.8%, PTX+siNFE2L2 = 61.2%) These findings implicate ELF3, FOSL1 and NFE2L2 in nuclear morphology maintenance and cytokinesis completion necessary for successful mitosis.

Through this model we aimed to assess how the combination of siRNA knockdown and paclitaxel synergized to disrupt the cell cycle and transitions between phases. For each individual or combination perturbation, we computed the inferred phase duration for G1, S/G2 and M phases using the model’s homotypic transition rates, which represent the fraction of cells that remain in the same phase through the timestep (**Figure 5E**). The inhibition of ELF3 alone strongly increased cell cycle duration (DMSO+siNonTarget = 49 hours, DMSO+siELF3 = 115 hours), with substantial increases to the time spent in G1 (DMSO+siNonTarget = 7.9 hours, DMSO+siELF3 = 33.5 hours) and S/G2 (DMSO+siNonTarget = 37.7 hours, DMSO+siELF3 = 72.1 hours) phases. The combination of paclitaxel treatment and siRNA knockdown resulted in the longest cell cycle durations for both the PTX+siELF3 (160 hours) and PTX+siIRF9 (159 hours) conditions compared to PTX+siNonTarget (84.4 hours). To further assess the therapeutic impact of siRNA knockdown and paclitaxel combination treatments, we used a Highest Single Agent (HSA) model to compare the inferred phase duration under combination paclitaxel + siRNA to the highest inferred phase duration for a single agent (either paclitaxel alone or siRNA alone, **Figure 5F**)[47]. In this approach a duration ratio less than 1 indicates antagonism (siRNA + paclitaxel results in shorter inferred phase duration than highest single agent), a duration ratio of 1 means there is no benefit of combination compared to highest single agent, and a duration ratio greater than 1 indicates a positive synergistic effect (siRNA + paclitaxel results in longer inferred phase duration than highest single agent). We found that although knockdown of NFE2L2 alone resulted in longer cell cycle phases, these changes were not particularly synergistic with paclitaxel treatment. In contrast, IRF9 knockdown resulted in increased duration ratios of all three phases compared to HSA (G1: 1.93, S/G2: 1.87, M: 1.8), while FOSL1 knockdown resulted in increased duration ratios for G1 and M phases (1.96, 2.19 ratios respectively). ELF3 knockdown showed the greatest synergy for the G1 phase, with a G1 duration ratio of 2.55, indicating that the combination of ELF3 knockdown with paclitaxel treatment strongly inhibits cell cycle progression out of G1.

Motivated by these ELF3 findings, we hypothesized that ELF3 expression may be predictive of overall survival in breast cancer. To that end we assessed the METABRIC[48] breast cancer cohort and used ELF3 expression to stratify patients into three categories: ‘high’ (Upper quartile of ELF3 expression), ‘mid’ (Inter quartile range of ELF3 expression), and ‘low’ (Lower quartile of ELF3 expression). We found that ELF3 expression was prognostic in both directions, with ELF3-high tumors having significantly shorter recurrence free survival (HR = 1.21, p value = 0.025, Cox Proportional Hazard) and ELF3-low tumors having a significantly longer overall survival (HR = 0.77, p value = 0.004, Cox Proportional Hazard) compared to the ELF3-mid tumors (**Figure 5G**). We then compared the gene expression between the ELF3-high and ELF3-low groups and found that the ELF3-high tumors were significantly enriched for MSigDB hallmarks related to cell cycle progression (HALLMARK_G2M_CHECKPOINT: NES = 1.99, Benjamini-Hochberg FDR = 5.0e-8, HALLMARK_E2F_TARGETS: NES = 1.94, Benjamini-Hochberg FDR = 2.9e-7, **Figure 5H, Supplemental Figure 8A**). We also found that the ELF3-low tumors were enriched for MSigDB hallmarks related to Allograft Rejection and Epithelial to Mesenchymal Transition (NES = - 2.14, Benjamini-Hochberg FDR = 4.1e-11, and NES = -1.95, Benjamini-Hochberg FDR = 6.1e-8 respectively). These results support our experimental *in vitro* findings that ELF3 activity contributes to continued malignant cell proliferation, and that high ELF3 expression in human breast cancer is associated with cell cycle progression and is also a negative predictor of progression free survival.

## DISCUSSION

Paclitaxel is a cornerstone therapy for TNBC and is an important component of first line neoadjuvant treatment for newly detected disease. Despite this, less than 20% of breast cancer patients treated with combination neoadjuvant therapy (paclitaxel followed by combination fluorouracil + doxorubicin + cyclophosphamide) achieve pathological complete response (pCR), and 47% of TNBC patients without pCR have recurrent disease within 10 years[49]. Although long-term chemotherapy resistance is often facilitated by clonal selection for growth-permissive mutations[50–52], newer molecular profiling techniques have revealed that short-term adaptive responses are possible through rapid epigenetic changes without acquisition of new mutations [53, 54]. In this study, we sought to identify adaptive responses that emerge after paclitaxel treatment and that may be targeted to deepen therapeutic response. To that end, we characterized the phenotypic and transcriptional responses of

TNBC cells to paclitaxel, with a focus on changes in cell number, multinucleation, and transcription factor programs Using siRNA knockdown, live-cell imaging, and computational modeling, we identified several TFs that phenocopied key aspects of paclitaxel response, including reduced proliferation rates and an increased proportion of multinucleated cells. ELF3 knockdown *in vitro* was synergistic with paclitaxel treatment and suppressed G1 to S/G2 cell cycle progression. Analysis of the METABRIC breast cancer cohort revealed that high expression of ELF3 was associated with worse outcome and higher cell-cycle related pathway activity. Together, these findings support the idea that upregulation and activation of ELF3 is an early and transcriptionally based mechanism of paclitaxel resistance in TNBC.

Many drug and gene manipulation studies focus primarily on viability or other cell count proxies at a terminal timepoint[55–58]. While such cell viability studies have proven valuable, more recent studies have demonstrated that chemotherapies modulate multiple cancer-associated hallmarks, including cell cycle phase behavior, senescence and nuclear morphology[42, 59, 60]. Further, there is evidence that the complex behavior of cellular systems are inherently dynamic, and their complex behaviors are better understood with measures that capture temporal behavior[43, 61–63]. While our live-cell studies captured important changes in cell cycle dynamics and the population distribution of various cell cycle states, no single metric captures the complete biological response. Future studies could deploy a richer panel of reporter molecules to gain deeper insights into other aspects of the response, including the timing and order of transcription factor activation, activation of specific cell cycle checkpoints, and activation of senescence or apoptotic pathways[64–66].

In this study we identified dual roles of the transcription factor ELF3 that contribute to paclitaxel tolerance by: 1) permitting cells to transition from G1 to S/G2, and 2) enabling successful division into two mononuclear daughter cells. These findings were enabled by a Markov Model of cell cycle progression built on population level cell count data which learned the transition rates between cell cycle phases and inferred cell cycle phase durations[42, 46]. While the inferred cell cycle durations represent an accurate prediction of the population’s average behavior, they cannot inform whether this arises from a homogenous or heterogenous distribution of cell cycle durations. This is of particular interest in the case of cancer treatment, as a small population of cells with a fitness advantage may eventually overtake the other populations, thus achieving therapeutic resistance[67]. An alternative approach could track individual cells and their progeny to build complete lineages with accompanying cell cycle timing information. Lineage based approaches tend to be relatively low throughput due to the computational and experimental requirements, but offer the opportunity to discern between heterogenous states of differing cycling speeds[46]. Another limitation of our Markov Model’s implementation is the assumption that transition rates are static throughout the duration of observations. While the output of the model mapped well within the 72-hour measurement window, there was some divergence at the end of the experiment that may suggest a weakening of either siRNA or paclitaxel effect. Incorporation of temporal information could be used to the current model implementation and could be useful for predicting combination drug effects and optimizing the drug schedule for maximum disruption of cell cycle progression.

Paclitaxel inhibits cell growth by simultaneously promoting microtubule assembly and inhibiting microtubule depolymerization, which results in mitotic checkpoint failure and subsequent apoptosis or senescent arrest[68, 69]. The *in vitro* experimentation performed in this study represents an extensive investigation into the phenotypic and molecular responses of TNBC cells to paclitaxel, however we acknowledge that tumors are comprised of diverse cell types and intercellular signaling molecules can influence therapeutic response in breast cancer and other malignancies[70–72]. Indeed, the tumor microenvironment is known to have a significant impact on drug response through cell-cell interaction and alterations to extracellular matrix[73, 74]. While we did not include stromal cells in our study, our findings of paclitaxel induced upregulation chemokines (CXCL1, CXCL8) support the idea that malignant cells that persist through paclitaxel treatment will have differential interactions with the immune system as compared to treatment-naïve cells. Tumor-derived CXCL1 is known to recruit immunosuppressive myeloid cells that inhibit CD8^+^ T cell infiltration[75]. The chemokine CXCL8 plays multiple pro-tumorigenic roles including recruitment of immunosuppressive neutrophils[76], promotion of angiogenesis[77] and maintenance of breast cancer stem cells[78]. Future studies that more deeply consider the influence of stromal and immune cells signals in modulating therapeutic response will be needed to better understand the complete system of factors involved in paclitaxel resistance.

As key regulators of multiple molecular programs, many transcription factors are known to contribute to cancer-associated phenotypes[79] and therapeutic response[80, 81]. Our study found that the ETS family transcription factor ELF3 was upregulated during early response to paclitaxel treatment, and siRNA knockdown of ELF3 was synergistic with paclitaxel treatment at slowing cell line growth. Other studies have found that high ELF3 activity is associated with inhibition of epithelial to mesenchymal transition [82]. Furthermore, inhibition of ELF3 was found to reduce proliferation across a number of cancer models including lung adenocarcinoma[83], neuroendocrine carcinoma[84] and prostate cancer[85]. Circulating tumor cells have elevated ELF3 expression in both murine models and human breast cancer[86]. Conserved dysregulation of ELF3 across cancer types may be related to its genomic location (loci 1q32) which is commonly amplified across cancers[87, 88] and also encodes for a number the cancer related genes including MDM4 (p53 suppressor)[89, 90].

Taken together, this work has identified ELF3 upregulation as an acquired mechanism of paclitaxel resistance. These findings support the development of pharmacological agents that inhibit ELF3 activity and could be used in combination with paclitaxel to further improve patient outcomes. While it has been historically difficult to develop targeted transcription factor inhibitors due to their lack of enzymatic activity, recent advances, such as targeted siRNA nanoparticles and indirect inhibition through targeting multiple interacting proteins, have made pharmacomodulation of transcription factors more tenable[91–93]. Until such therapies are developed, ELF3 may serve as a useful biomarker which predicts the development of paclitaxel resistance and continued malignant proliferation.

## METHODS

### Cell culture

HCC1143 (ATCC), HCC1806 and MDA-MB-468 cells were authenticated by STR profiling and tested negative for mycoplasma. HCC1143 and HCC1806 cells were cultured in RPMI 1640 with L-glutamine (cat. 11875119, Life Technologies Inc.) supplemented with 10% fetal bovine serum (#16000-044, Gibco). MDA-MB-468 cells were cultured in DMEM (#11965-092, Life Technologies Inc.) supplemented with 10% fetal bovine serum (#16000-044, Gibco). All lines were incubated at 37C with 5% CO_2_. For perturbation experiments, cells were seeded into appropriate assay vessel for 24 hours prior to treatment with either vehicle control (DMSO; PBS) or perturbation (table below).

| <u>Perturbation</u> | <u>Shorthand</u> | <u>Concentration used</u><br><u>for scRNA-seq</u> | <u>Source</u> | <u>Identifier</u> | <u>Vehicle</u> |
| --- | --- | --- | --- | --- | --- |
| Paclitaxel | PTX | 1nM | LC Labs | P-9600 | 0.1% DMSO |
| Notch Inhibitor | NOTCHi | 1uM | Millipore Sigma | BM0018-5MG | 0.1% PBS |
| Interferon Beta | IFNB | 20ng/mL | PBL Assay<br>Science | 11410-2 | 0.1% PBS |
| Interferon Gamma | IFNG | 20ng/mL | R&D Systems | 385-IR-100 | 0.1% PBS |
| Transforming<br>Growth Factor Beta | TGFB | 10ng/mL | Biotechne | 7754BH005 | 0.1% PBS |
| Lymphotoxin | LT | 10ng/mL | Biotechne | 8884-LY-025 | 0.1% PBS |
| Oncostatin M | OSM | 10ng/mL | Cell Signaling<br>Technology | 5367SC | 0.1% PBS |

### Fixed cell assays

Cells were plated at 3000 cells in 100ul of complete media per well in a 96 well plate (#08-772-225, FisherScientific). After 24 hours, an additional 100ul of either vehicle (0.1% DMSO) or paclitaxel containing complete media was added. After 72 hours cells were fixed with 4% Formaldehyde (#28908, ThermoFisher Scientific) for 15 minutes at room temperature, then permeabilized with 0.3% Triton X-100 (#X100-100ML, Sigma Aldrich) for 10 minutes at room temperature, then washed twice with PBS. Fixed cells were blocked with 1% BSA (A7906-100G, Millipore Sigma) in PBS for 1 hour at room temperature and then stained overnight with 1:100 anti-CDKN2A/p16INK4A+CDKN2B/p15INK4B-AF644 (#ab199756, Abcam), and 1:100 anti-cPARP-AF647 (#6987S, Cell Signaling Technology) or 1:500 anti-TUBB3-AF647 (#ab190575, Abcam) overnight at 4C. Each well was washed twice with room temp PBS then stained with 0.5ug/mL DAPI (4083S, Cell Signaling Technology) in PBS for 15 minutes at room temperature. Following DAPI staining, wells were washed once with PBS, then stained with 1:20,000 HCS CellMask in PBS (Orange: #H32713, Green: #H32714, Invitrogen) for 15 minutes at room temperature. Wells were washed twice with room temperature PBS and then 4 fields of view per well imaged on an InCell 6000 (GE Healthcare). Images were segmented with two custom Cellpose[45] models to segment the nucleus (from DAPI channel) and cytoplasm (from HCS Cellmask channel). Image quantification was performed in R (v4.3.1) using EBImage (v4.42.0), and cells were annotated based on the number of distinct nuclei segmented within each cytoplasmic mask.

### scRNA-seq library preparation and sequencing

Experiment 1 (DMSO 24 hour, DMSO 72 hour, Paclitaxel 24 hour, Paclitaxel 72 hour): Each condition had a single-cell RNA library prepared using 10X Genomics Single Cell 3’ v2 kits and sequenced on an Illumina NextSeq 500 for 500e6 reads per library.

Experiment 2: All conditions were multiplexed using Hashtag Oligonucleotide barcoding technology (TotalSeq-B, Biolegend) following manufacturer standard protocol. A paired feature-barcode library and mRNA library were generated using the Single Cell 3’ v3 kit (10X Genomics) following manufacturer instructions and then sequenced on an Illumina NovaSeq for 800e6 reads.

### scRNA-seq data processing

For both experiments; raw base call files were converted to FASTQ format with bcl2fastq (Illumina). Cellranger count (v6.0.2) was used to align reads to the GRCh38 transcriptome (GRCh38-2020-A, accessed from 10X Genomics) and count UMI reads. The R package Seurat[94, 95] (4.0.5) was used to perform variable feature identification, linear and nonlinear dimensionality reduction, unsupervised clustering and differential gene expression.

Variance Stabilizing Transformation was used to identify the top 2000 variable genes and Principal Component Analysis (PCA) was used to reduce these 2000 genes to 10 components for UMAP embedding and unsupervised clustering. Differential expression analysis was performed using the FindMarkers function of Seurat with default parameters. Geneset enrichment analysis was performed with the R package clusterProfiler[96] (v4.8.2) using significantly upregulated genes compared to time-matched vehicle control (abs(avg_log2FC) > 0.5, Benjamini Hochberg FDR < 0.05).

### Transcription Factor Enrichment Analysis

Significantly upregulated genes (avg_log2FC > 0.5, Benjamini Hochberg FDR < 0.05) were computed for paclitaxel, IFNB and IFNG treated samples compared to time-matched vehicle treated cells. ChEA3 enrichment analysis was performed with default settings using R code from the CHEA3 API documentation (https://maayanlab.cloud/chea3/) to perform an online query using either the genes uniquely upregulated in paclitaxel treated cells, or those shared between paclitaxel and either of the interferon responses. The top 15 ranked transcription factors from both the paclitaxel unique and paclitaxel-interferon shared TF enrichment lists were considered when nominating siRNA knockdown targets. Any TF that also had at least 0.25 log2 fold change for paclitaxel at either 24 or 72 hours compared to vehicle control was included in the siRNA knockdown panel.

### siRNA Knockdown

Cells were plated in 90ul of serum free media per well of a 96 well plate. 24 hours later, siRNA knockdown mixture was prepared using a cell-line optimized concentration of Lipofectamine RNAiMAX (cat 13778075-075, Invitrogen) and siRNA (Horizon Discovery ON-TARGETplus) following RNAiMAX recommended protocol. The final concentration of siRNA per well was 1pmol and the final volume of RNAiMAX per well was 75nL for HCC1143, and 37.5nL for HCC1806 or MDA-MB-468 in 100uL of cell containing volume. 24 hours after siRNA transfection cells were treated with an addition of 100uL complete media containing either DMSO vehicle control or paclitaxel.

### Protein isolation

Protein isolation: siRNA knockdown of HCC1143 cells was performed using the siNonTarget, siELF3, siFOSL1, and siNFE2L2 pools as described above. After 24 hours of knockdown, perturbation containing media was added such that media volume doubled and had a final concentration of either 0.1% DMSO (vehicle control) or 1nM Paclitaxel. After 72 hours of perturbation, cells were washed with 4C PBS then lysed by 5 minute incubation at 4C with RIPA buffer (R0278, Sigma) supplemented with 1X Halt Protease and Phosphatase Inhibitor Cocktail (1861281, Thermo Scientific). Remaining cells were scraped from the plate and lysate was snap frozen in liquid nitrogen then stored at -80C overnight. The following day lysate was clarified by centrifugation at 21,130 x g for 10 minutes at 4C. The supernatant was collected and the protein concentration was immediately quantified. Remaining protein was stored at -80C.

### Western Blot

Protein quantification was performed using the Western Simple protocol on the Jess capillary western machine using the 12-230 kDa cartridge and following manufacturer instructions (Biotechne). Primary antibodies targeting the protein products of ELF3 (anti-ESE1, ab133521, Abcam), FOSL1 (anti-FRA1, sc28310, Santa Cruz), and NFE2L2 (anti-NRF2, HPA043438-1, Sigma) were used at 1:50 dilution. Lysates were loaded at a concentration of 2mg/mL and volume of 5uL per capillary well, and the Anti-rabbit detection kit (DM-001, Biotechne). was used to quantify primary antibody levels. Peak quantification was performed using the included Compasssoftware with default settings (v6.3.0, Biotechne).

### HDHB reporter live-cell assays

siRNA knockdown and drug treatment was performed as described above, and then the plate was loaded on an Incucyte S3 (Sartorious) and cells imaged every 15 minutes for 72 hours post drug treatment. At each timepoint 4 fields of view were captured at 20x magnificantion in each well using the phase, red and green channels. A cytoplasmic mask was computed from the mean of normalized red/green channel, and a nuclear mask was computed from the red channel using custom trained Cellpose[45] models. Image quantification was performed in R (v4.3.1) using EBImage (v4.42.0). An additional perinuclear ring mask was computed as the 11 pixel dilation from the nuclear mask, but still bound by the cytoplasmic mask. To determine mClover localization thresholds for cell cycle assignment, 250 cell images were randomly selected and manually assigned to the G1, S/G2 or M cell cycle state based on mClover localization. The mClover intensity ratios were then used to determine thresholds for automated cell cycle phase calling which was applied to the rest of the data set (**Supplemental Figure 5A**). Mononuclear cells with a Perinuclear:Nuclear mean intensity ratio greater than 0.8 and Nuclear:Cytoplasmic total intensity less than 0.5 were assigned to the S/G2 phase. Mononuclear and Multinuclear cells with a Nuclear:Cytoplasmic total intensity ratio greater than 0.8 and Perinuclear:Nuclear mean intensity ratio less than 0.8 were assigned to the ‘M’ phase. The remainder of mononuclear cells were assigned ‘G1’, and the remainder of multinucleated cells were assigned ‘Multinucleated’.

### Markov modeling

The 5-frame moving average of cell count per cell cycle phase was downsampled to one value per hour and used to train a markov model for each unique siRNA (NonTarget, ELF3, FOSL1, NFE2L2, IRF9, PLK1) +/- paclitaxel condition. The transition matrix of the model was constrained such that cells could remain in their current phase, progress through the cell cycle (G1 -> S/G2, S/G2 -> M, M -> G1 with replication) or transition from M phase to an absorbing (permanent) multinucleated phase. Models were trained for 15 epochs, and the first epoch was seeded with an identity transition matrix. 3000 random transition matrices were generated each epoch, and the 5 with lowest error were used as seeds for the following epoch. The prior best performing matrices were updated with randomly generated matrices at a learning rate of 0.1 for the first epoch, halving every 2 epochs.

The prediction for counts for each future state (S_n+1_) is calculated as the product of the counts at the prior state (S_n_) by the transition matrix (P) and the replication matrix (RM).

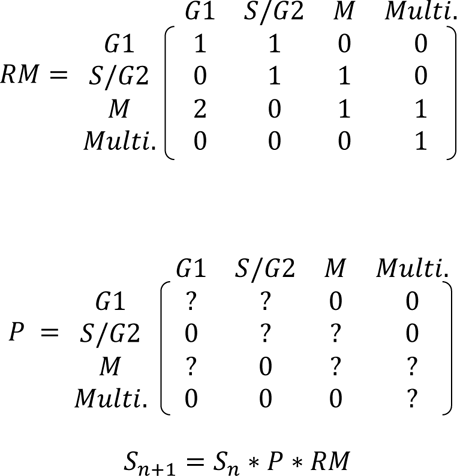

The error of the markov predicted cell counts (c_exp_) compared to observed counts (c_obs_) was computed as the arithmetic mean of the Root Mean Squared Relative Error (RMSRE) of each cell cycle phase across all predicted timepoints. The noise floor of RMSRE was estimated with a second-order loess fit with span of 0.75 (loess function from R package ‘stats’, v4.3.1).

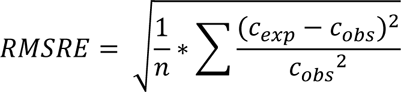

The mitotic success rate (MSR) of each condition was computed as the ratio of M-to-G1 transition (P_M,G1_) to the sum of the transition rates for M-to-G1 (P_M,G1_) and M-to-multinucleated (P_M,Multi_):

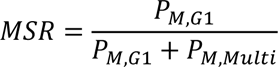

The expected duration of G1, S/G2 and M cell cycle phases was calculated from the homotypic transition rates as[97]:

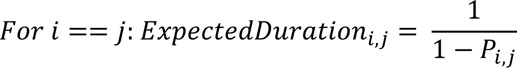

### Metabric survival and microarray analysis

The Metabric[48] microarray and patient metadata was accessed through cbioportal[98–100] and analyzed using R (v4.3.2) and the ‘survival’ package (v3.5.7). The z-scored microarray expression data was used to categorize patients into ‘high’ (highest expressing quartile), ‘mid’ (first to third expressing quartile) or ‘low’ (lowest expressing quartile) based on expression of ELF3. For survival analysis, patients were filtered to those with microarray data and then Kaplan-meier survival curves were generated with the ‘ggsurvfit’ package (v1.0.0). Cox proportional hazard statistics were calculated with the ‘coxph’ function of the ‘survival’ package (v3.5.7). Differential expression was calculated from the log normalized microarray data using the ‘wilcoxauc’ function from the ‘presto’ package (v1.0.0). Significantly differentially expressed genes (abs(logFC) > 0.5 and adjusted p < 0.05) where used to compute MSigDB hallmark GSEA using the ‘clusterprofiler’ (v4.10.1) and ‘msigdbr’ (v7.5.1) packages.

Full reagent list:

| reagent | shorthand | type | source | identifier |
| --- | --- | --- | --- | --- |
| HCC1143 | - | cell line | ATCC | CRL-2321 |
| HCC1806 | - | cell line | ATCC | CRL-2335 |
| MDA-MB-468 | - | cell line | ATCC | HTB-132 |
| RPMI 1640 | RPMI | reagent | Life Technologies | 11875119 |
| DMEM | DMEM | reagent | Life Technologies | 11965-092 |
| Fetal Bovine Serum | FBS | reagent | Gibco | 16000-044 |
| Dimethyl Sulfoxide | DMSO | reagent | Millipore Sigma | D8418-250ML |
| Phosphate Buffered Saline | PBS | reagent | Gibco | 14190235 |
| Paclitaxel | PTX | reagent | LC Labs | P-9600 |
| Interferon-Beta | IFNB | reagent | PBL Assay Science | 11410-2 |
| Interferon-Gamma | IFNG | reagent | R&D Systems | 385-IR-100 |
| Human Oncostatin M | OSM | reagent | Cell Signaling Technology | 5367SC |
| Recombinant Human Lymphotoxin alpha1/beta2 protein | LT | reagent | Biotechne | 8884-LY-025 |
| Recombinant Human TGFB-Beta 1 | TGFB | reagent | Biotechne | 7754BH005 |
| BMS-906024 | NOTCHi | reagent | Millipore Sigma | BM0018-5MG |
| 16% Formaldehyde (w/v) | - | reagent | ThermoFisher Scientific | 28908 |
| Triton X-100 | - | reagent | Millipore Sigma | X100-100ML |
| Normal Goat Serum Blocking Solution | - | reagent | MP Biomedicals | #0219135680 |
| Lipofectamine RNAiMAX transfection Reagent | - | reagent | ThermoFisher Scientific | 13778075 |
| Bovine Serum Albumin | BSA | reagent | Millipore Sigma | A7906-100G |
| RIPA buffer | - | reagent | Sigma | P0278 |
| 100X Halt Protease and Phosphatase Inhibitor Cocktail | - | reagent | ThermoFisher Scientific | 1861281 |
| anti-CDKN2A/p16INK4A+CDKN2B/p15INK4B-AF644 | p16 | antibody | Abcam | ab199756 |
| anti-cPARP-AF647 | cPARP | antibody | Cell Signaling Technology | 6987S |
| anti-TUBB3-AF647 | TUBB3 | antibody | Abcam | ab190575 |
| HCS CellMask Green | CellMask Green | stain | Invitrogen | H32713 |
| HCS CellMask Orange | CellMask Orange | stain | Invitrogen | H32714 |
| DAPI | DAPI | stain | Cell Signaling Technology | 4083S |
| anti-ESE1 | - | antibody | Abcam | ab133621 |
| Fra1 Antibody (C-12) | - | antibody | Santa Cruz | sc-28310 |
| Anti-NFE2L2 antibody produced in rabbit | - | antibody | Millipore Sigma | HPA043438 |
| siATF3 Smartpool | siATF3 | siRNA | Hoizon Discovery | L-008663-00 |
| siDDIT3 Smartpool | siDDIT3 | siRNA | Hoizon Discovery | L-004819-00 |
| siELF3 Smartpool | siELF3 | siRNA | Hoizon Discovery | L-016080-00 |
| siFOSL1 Smartpool | siFOSL1 | siRNA | Hoizon Discovery | L-004341-00 |
| siIRF7 Smartpool | siIRF7 | siRNA | Hoizon Discovery | L-011810-00 |
| siIRF9 Smartpool | siIRF9 | siRNA | Hoizon Discovery | L-020858-00 |
| siJUNB Smartpool | siJUNB | siRNA | Hoizon Discovery | L-003269-00 |
| siJUN Smartpool | siJUN | siRNA | Hoizon Discovery | L-003268-00 |
| siKIF11 Smartpool | siKIF11 | siRNA | Hoizon Discovery | L-003317-00 |
| siKLF6 Smartpool | siKLF6 | siRNA | Hoizon Discovery | L-021441-00 |
| siMAFF Smartpool | siMAFF | siRNA | Hoizon Discovery | L-003903-00 |
| siNFE2L2 Smartpool | siNFE2L2 | siRNA | Hoizon Discovery | L-003755-00 |
| siPLK1 Smartpool | siPLK1 | siRNA | Hoizon Discovery | L-003290-00 |
| siPLSCR1 Smartpool | siPLSCR1 | siRNA | Hoizon Discovery | L-003729-00 |
| siSP100 Smartpool | siSP100 | siRNA | Hoizon Discovery | L-015307-00 |
| ON-TARGETplus Non-targeting Control | siNonTarget | siRNA | Hoizon Discovery | D-001810-10 |
| ON-TARGETplus GAPD Control | siGAPD | siRNA | Hoizon Discovery | D-001830-10 |

## Data Availability

Single Cell RNA-seq data is available on the Gene Expression Omnibus with study ID: GSE266934 (https://www.ncbi.nlm.nih.gov/geo/query/acc.cgi?acc=GSE266934). The raw images and processed data from immunofluorescent stained HCC1143 are available on Zenodo: (doi: 10.5281/zenodo.11237850). The raw images and processed data from the siRNA panel, and the processed data from live-cell imaging study are available on Zenodo (doi: 10.5281/zenodo.11238552). The raw images from the live-cell experiments are available upon request.

## Code Availability

All code related to data processing and figure generation are available on Github (https://github.com/HeiserLab/PTX_manuscript).

## Contributions

N.L.C., T.A.L., Z.H., S.M.G. and L.M.H. designed experiments. N.L.C., T.L. and Z.H. performed experiments. N.L.C., H.Z. and M.D. performed data analysis. N.L.C. and L.M.H. wrote the manuscript with input from all authors.

## Acknowledgements

We thank Drs. Tahereh Ziglari and Ferdinando Pucci for assistance with proteomic validation of siRNA knockdown. OHSU Massively Parallel Sequencing Shared Resource receives support from the OHSU Knight Cancer Institute NCI Cancer Center Support Grant P30CA069533. This work was supported by the Jayne Koskinas Ted Giovanis Foundation for Health and Policy and NIH research grant U54-CA209988 (L.M.H.). N.L.C. acknowledges funding from the NCI/NIH under award number T32CA254888. The Jayne Koskinas Ted Giovanis Foundation for Health and Policy is a private foundation committed to critical funding of cancer research. The opinions, findings, conclusions or recommendations expressed in this material are those of the author(s) and not necessarily those of the Jayne Koskinas Ted Giovanis Foundation for Health and Policy or its respective directors, officers, or staff.

---

**Supplementary Figure 1:**
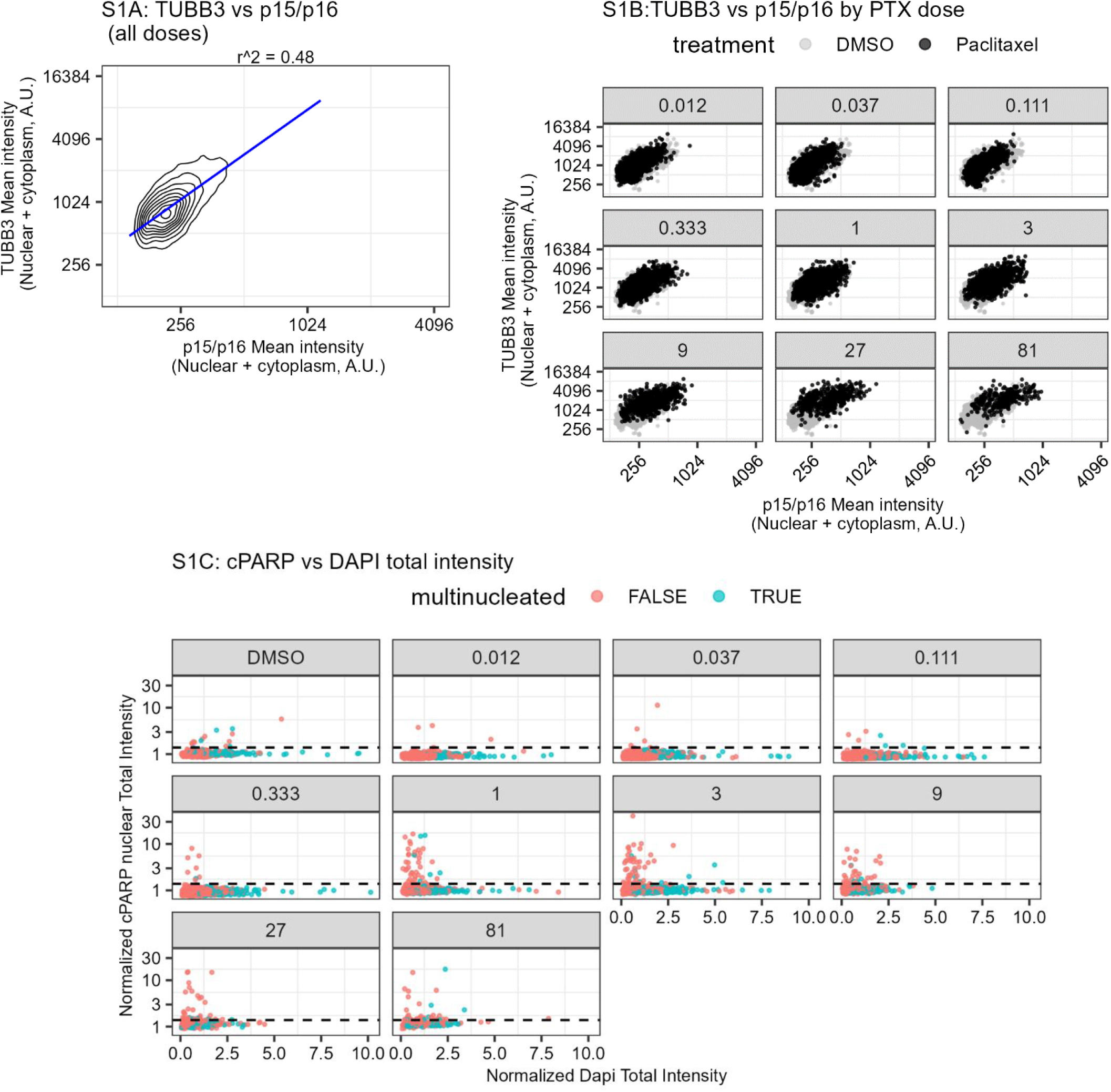
**S1A)** Density plot showing the distribution of cells for all conditions (DMSO + PTX). X axis shows mean intensity for p15/p16, y axis shows Mean Intensity for TUBB3 in arbitrary units (A.U.). R^2 squared shown for Pearson correlation (p-value < 2.2e-16). **S1B)** Breakout plots showing the same information as S1A. Control (DMSO) shown for every inset plot, and paclitaxel (PTX) for the nM dose listed above. **S1C)** Breakout plots showing the Normalized DAPI total intensity versus Normalized cPARP nuclear total intensity. Horizontal line indicates threshold for calling a cell ‘cPARP positive’, color indicates whether the cell is mononucleated (red) or multinucleated (blue).

**Supplementary Figure 2:**
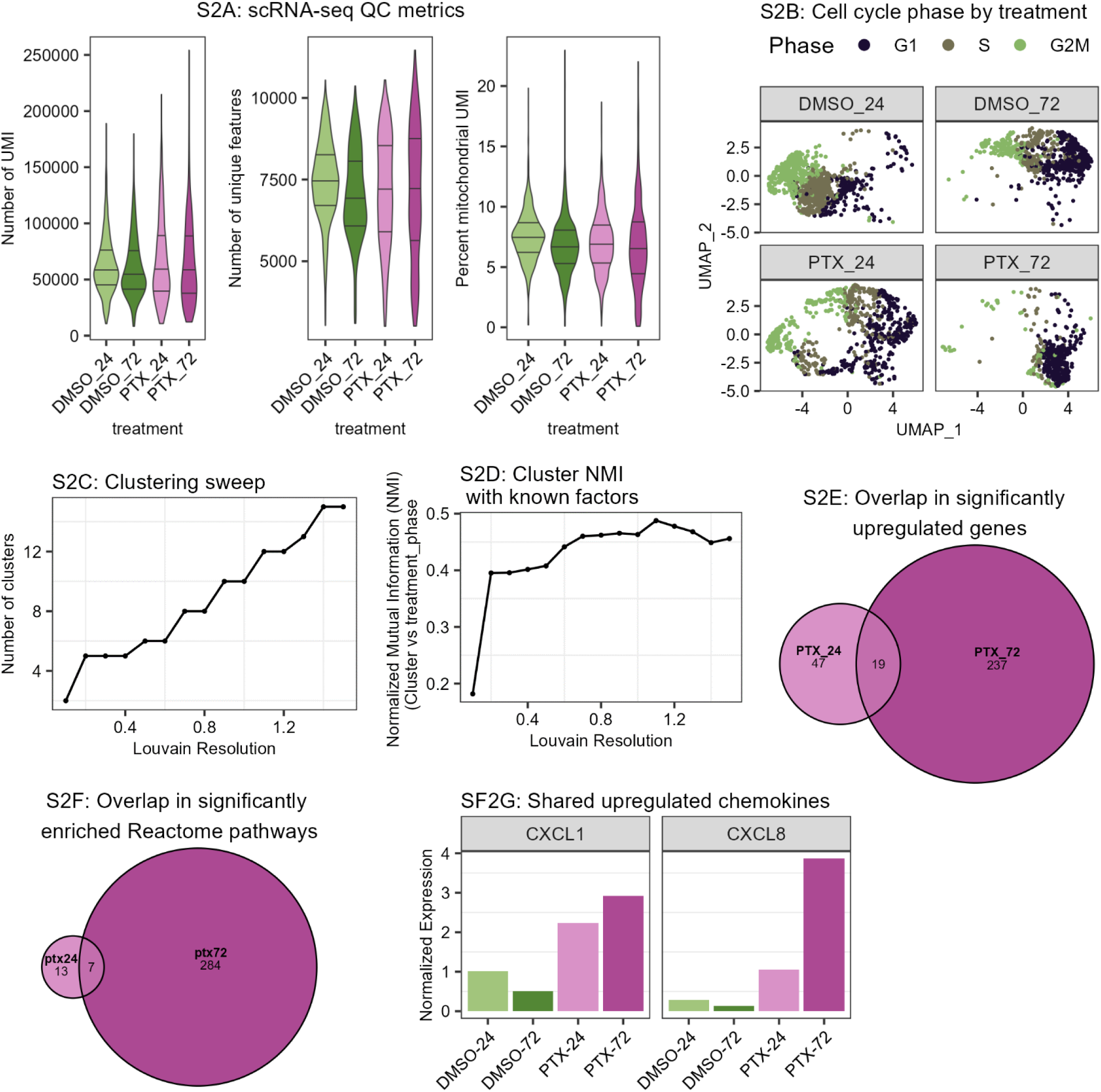
**S2A)** Violin plots of scRNA-seq QC metrics for the four conditions. Horizontal lines indicate the first, second and third quartiles. **S2B)** Breakout plots showing the same UMAP as Figure 2A split by condition and color coded by cell cycle phase. **S2C)** Number of clusters computed from Louvain clustering applied across a sweep of resolutions. **S2D)** The Normalized Mutual Information (NMI) between cluster label and biological label (treatment x cell cycle phase) computed across a sweep of Louvain clustering resolutions. The NMI values indicate that there is a high degree in overlap of information between unsupervised cluster labels and known biological labels. **S2E,F)** Euler plot showing the overlap in significantly upregulated genes (SF2E) and enriched Reactome pathways (SF2F) between paclitaxel at 24 hours (PTX24) and 72 hours (PTX72) compared to time matched control. **F2G)** Barplots showing mean expression of chemokines CXCL1 and CXCL8 which were significantly upregulated in both paclitaxel conditions compared to time matched control.

**Supplementary Figure 3:**
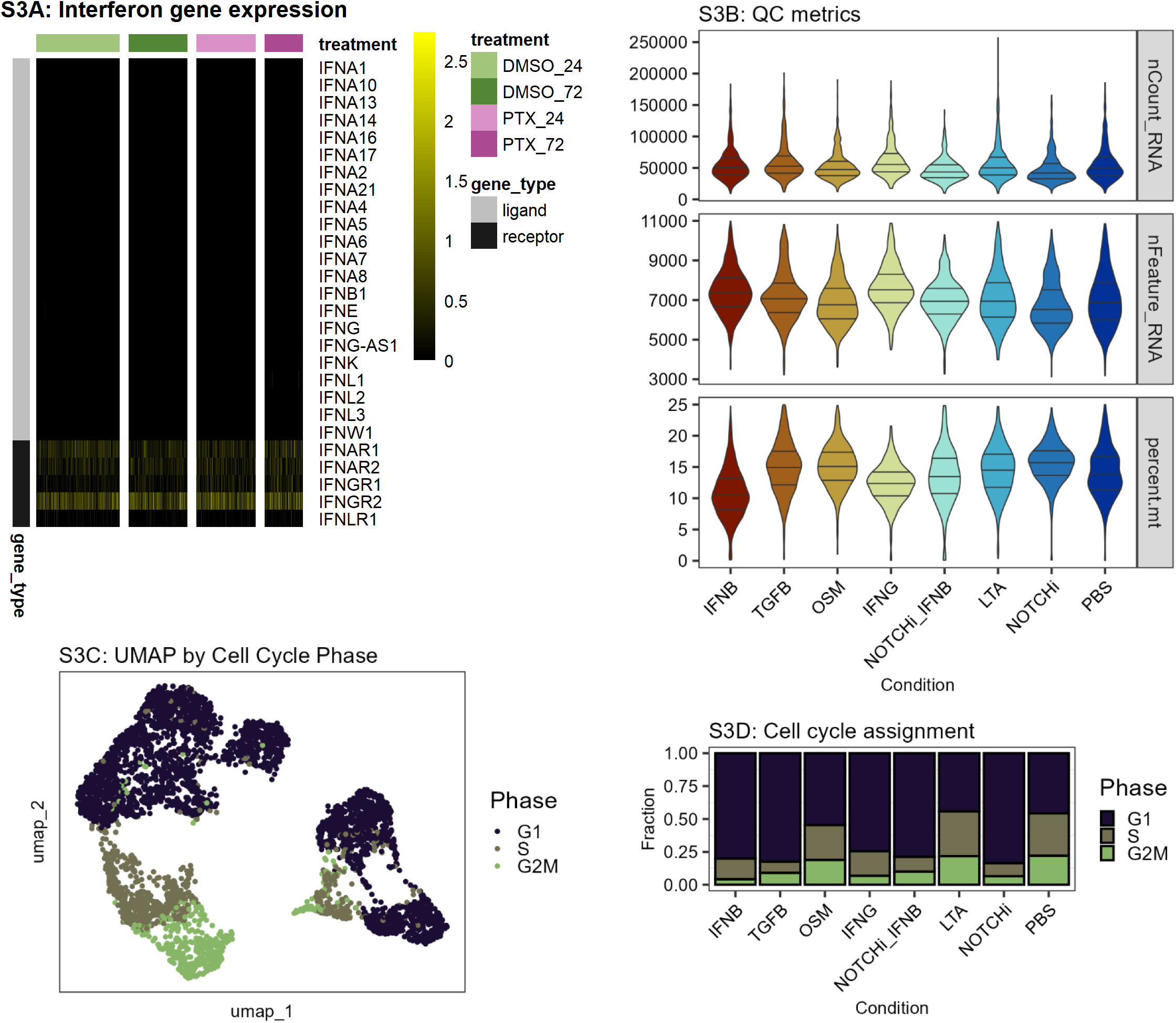
**S3A)** Heatmap showing expression for interferon ligands (gray) and receptors (black) for each of the paclitaxel scRNA-seq conditions. **S3B)** Violin plots of scRNA-seq QC metrics for the ligand perturbation conditions. Horizontal lines indicate the first, second and third quartiles. **S3C)** The same ligand perturbation scRNA-seq UMAP as Figure 3A, but color coded by cell cycle phase assignment. **S3D)** Bar plot indicating the proportion of cells assigned to each cell cycle phase for each condition.

**Supplementary Figure 4:**
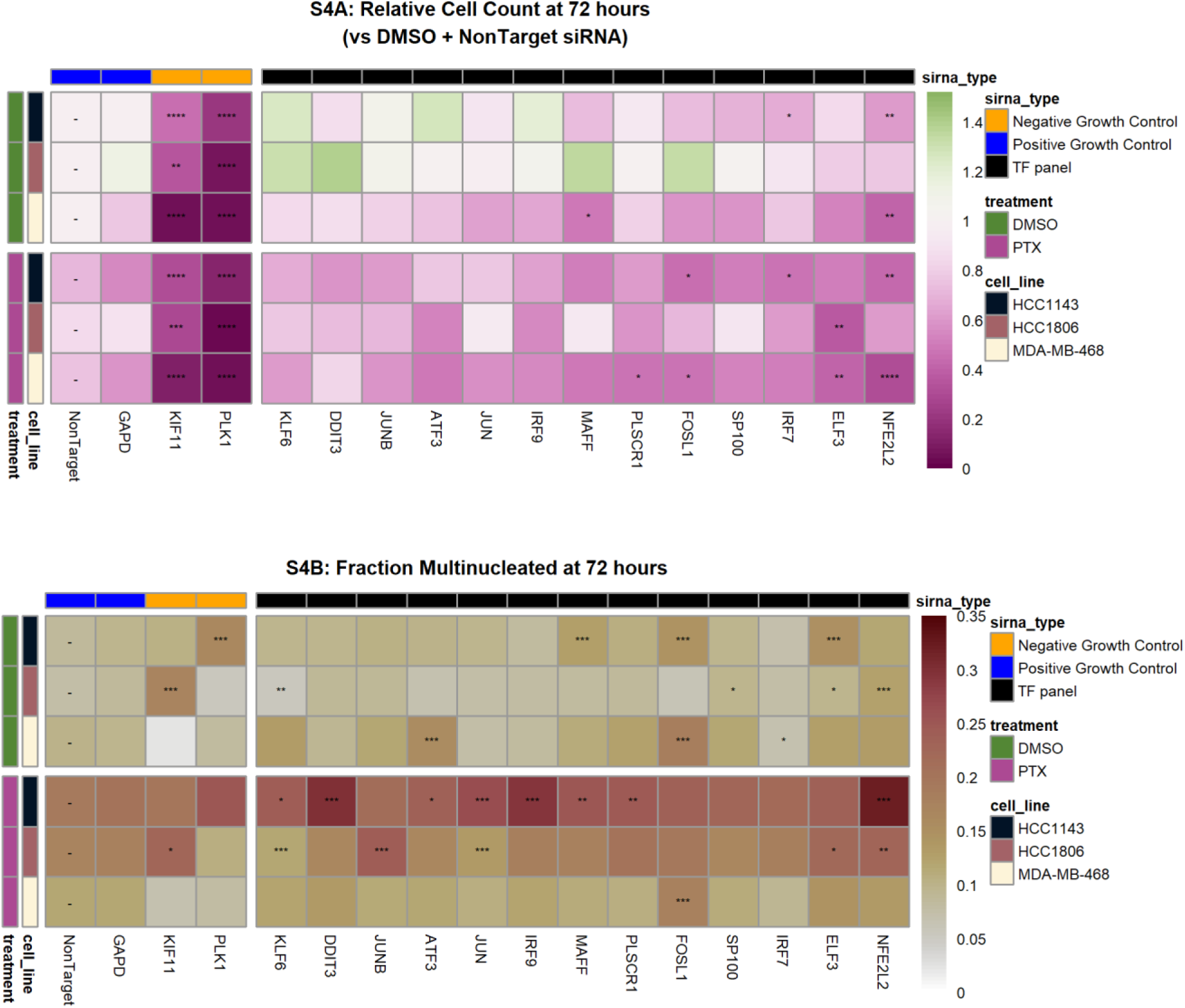
**S4A)** Heatmap of relative cell count (Normalized to NonTarget) for each siRNA +/- 1nM Paclitaxel (PTX) condition for the three Triple Negative Breast Cancer cell lines tested. GAPD and NonTarget siRNA are positive growth controls, KIF11 and PLK1 are negative growth controls. Significance assessed via t-test versus the same-drug (DMSO or PTX) NonTarget condition with Bonferroni Correction (p-values: *: < 0.05, **: < 0.01, ***: < 0.001). **S4B)** Heatmap representing the proportion of multinucleated cells for each siRNA +/- 1nM Paclitaxel (PTX) condition for the three Triple Negative Breast Cancer cell lines tested. GAPD and NonTarget siRNA are positive growth controls, KIF11 and PLK1 are negative growth controls. Significance assessed via proportions test versus the same-drug (DMSO or PTX) NonTarget condition with Bonferroni Correction (p-values: *: < 0.05, **: < 0.01, ***: < 0.001).

**Supplementary Figure 5:**
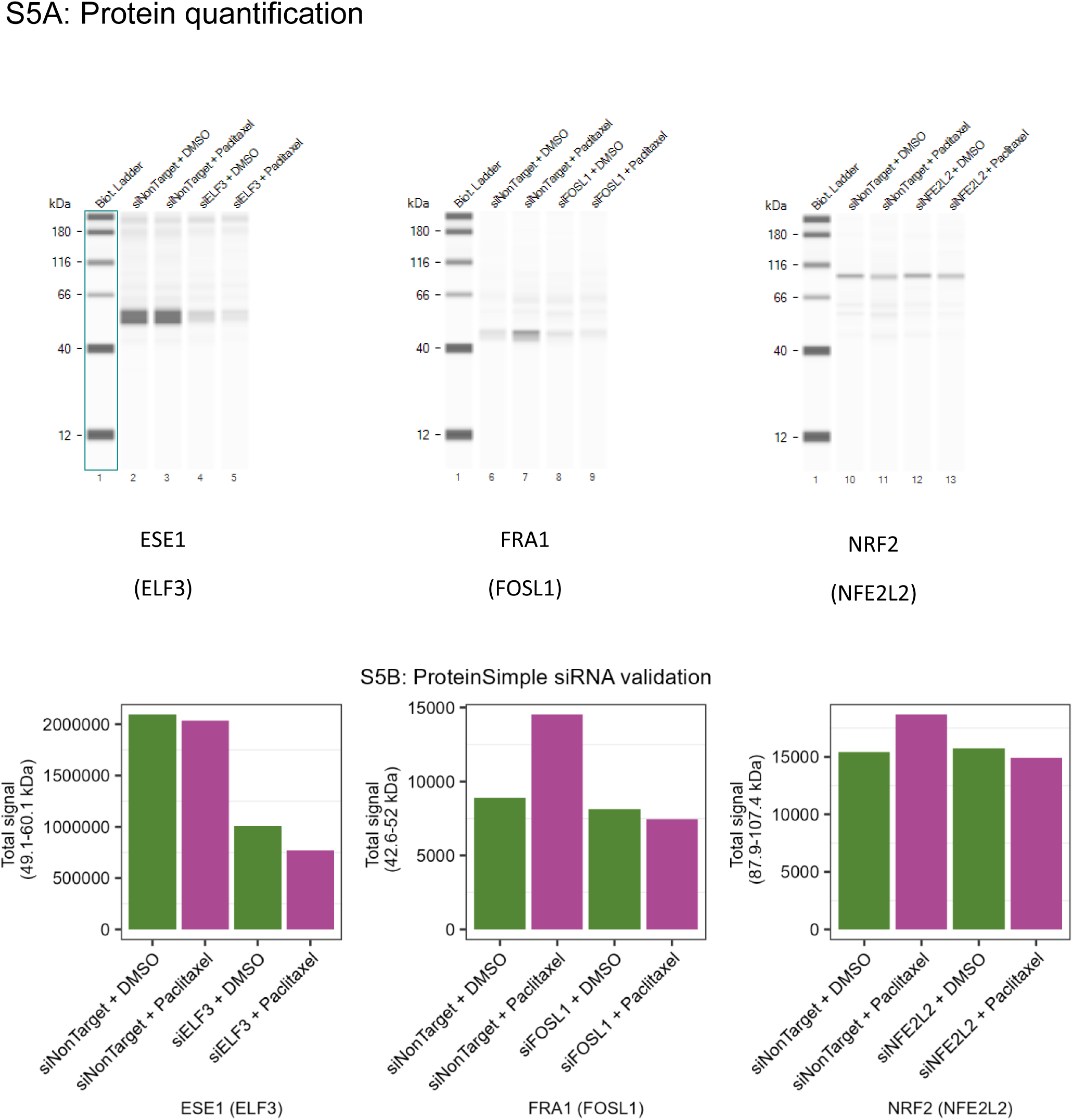
**S5A)** siRNA knockdown validation blots showing spectra images from ProteinSimple/WesternSimple protocol for ESE1 (ELF3), FRA1 (FOSL1) and NRF2 (NFE2L2) knockdown after 72 hours of treatment with either 0.1% DMSO or 1nM Paclitaxel. **S5B)** Quantification of the images above. Total signal indicates the sum of peak area for the +/- 10% range around the highest intensity peak.

**Supplementary Figure 6:**
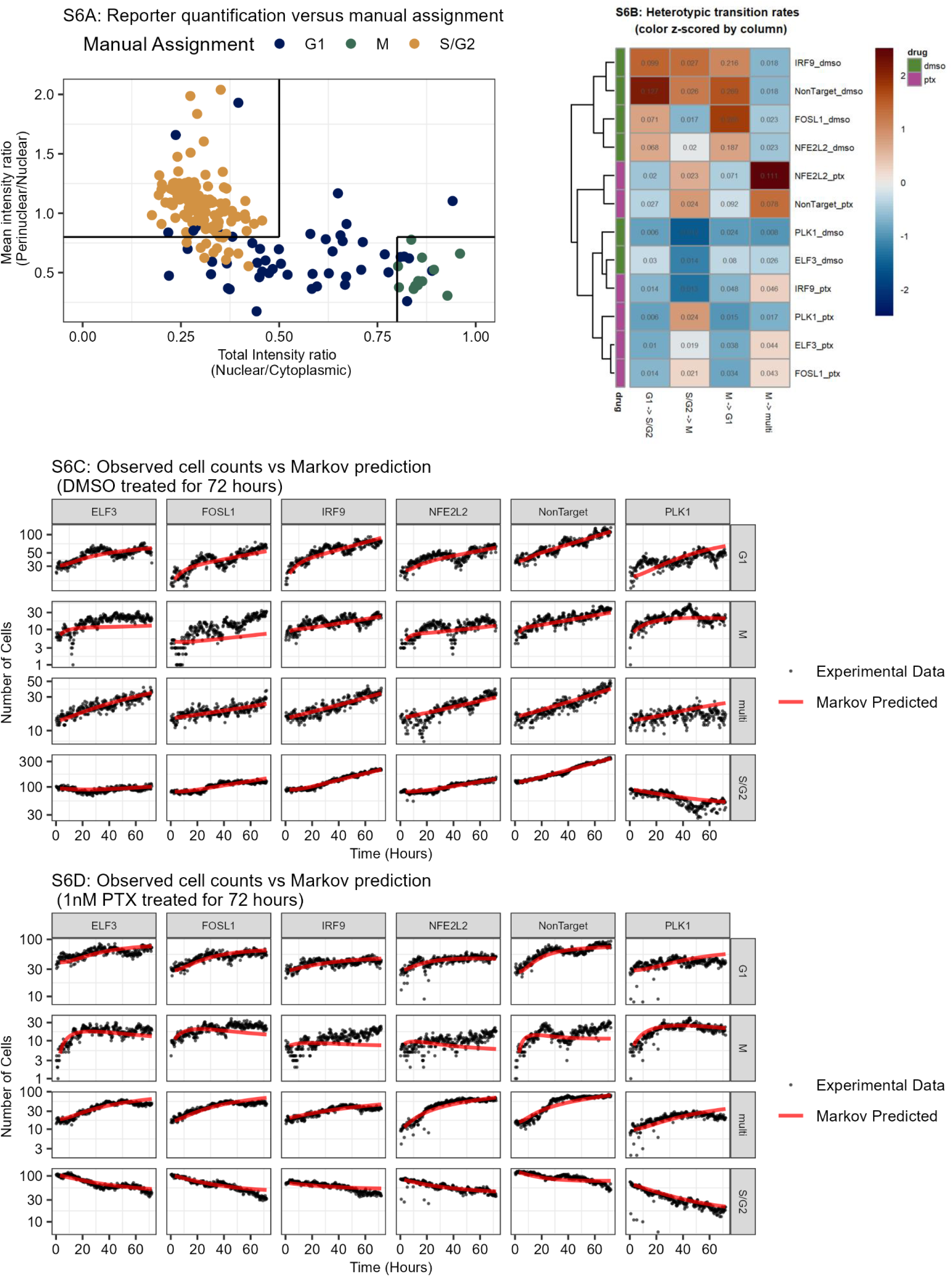
**S6A)** HDHB-mClover reporter intensities plot colored by manual assignment. 250 images of cells were randomly selected and manually assigned a cell cycle state (G1, M, S/G2) based on cell morphology and mClover intensity. The manual assignment was used to select Total Intensity Ratio (Nuclear vs cytoplasmic) and Mean intensity ratio (Perinuclear versus Nuclear) as defining features for automatic cell cycle assignment. Black lines represent thresholds used for automated cell cycle assignment. **S6B)** Heatmap showing the heterotypic (between different states) transition rates learned by the Markov model for each unique siRNA +/- Paclitaxel (PTX) condition. Inset number is the transition rate and color is the z-score of row. **S6C, SF6D)** Cell count over time plots for each of the DMSO (S6C) and PTX (S6D) treated conditions showing the experimental data (black dots) and Markov values (red line) predicted using the learned transition rates and initial time point.

**Supplementary Figure 7:**
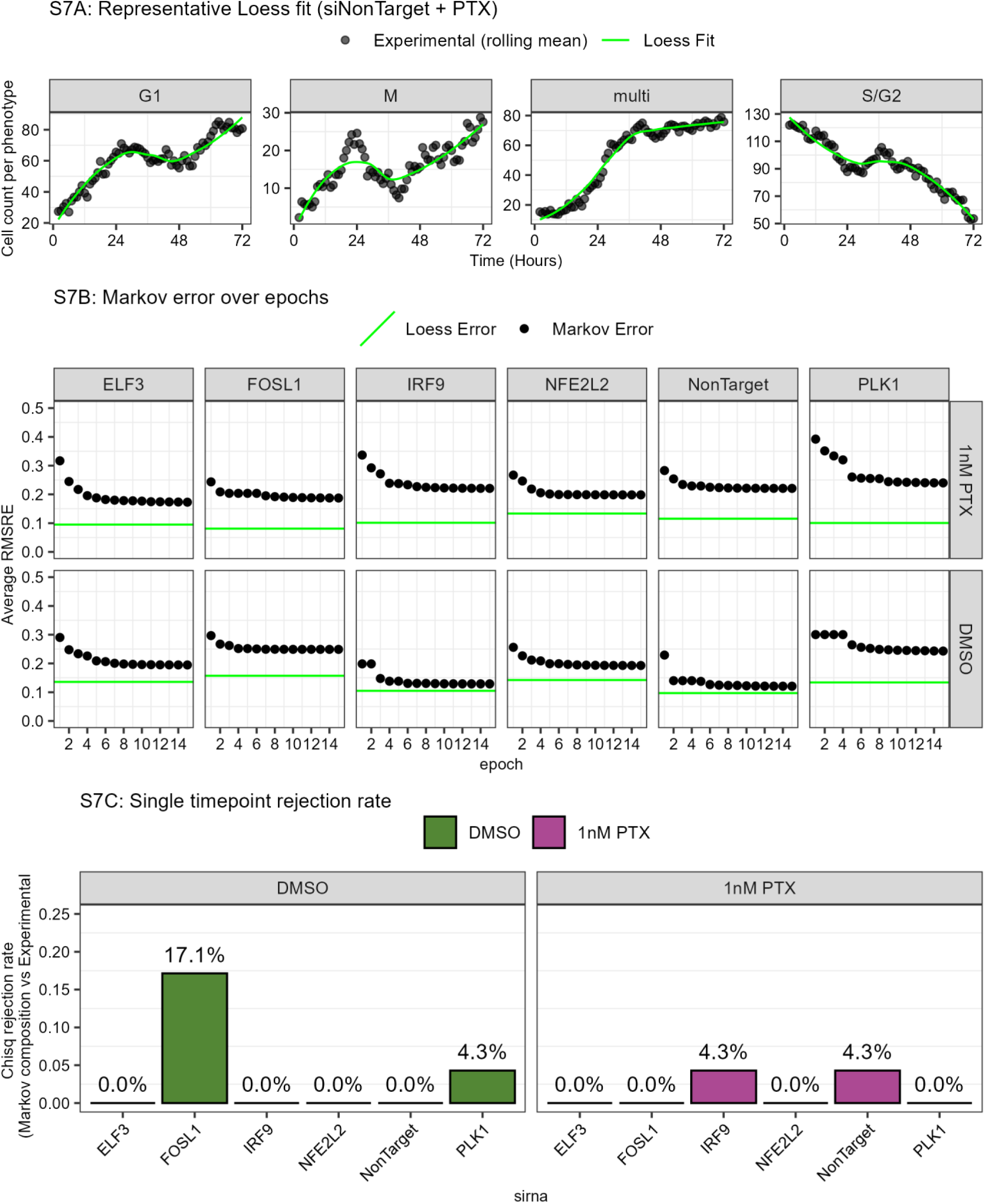
**S7A)** Representative plot showing the smoothed experimental counts (5-timepoint rolling mean) versus a Loess fit for the siNonTarget + Paclitaxel condition. **S7B)** Dot plots showing the Root Mean Squared Relative Error (RMSRE) for the Markov Model (black dots) over each training epoch versus the Loess fit (green line). Loess fit represents an estimate of the ‘noise floor’ of the measurement. **S7C)** Model rejection rate for each condition computed from the Chi-squared test applied between the experimental and model predicted composition for each single time point. A nominal Chi-squared p value < 0.05 was considered a significantly different timepoint. A rejection rate of 0% means that there was no significant difference in phenotype composition at any timepoint.

**Supplementary Figure 8:**
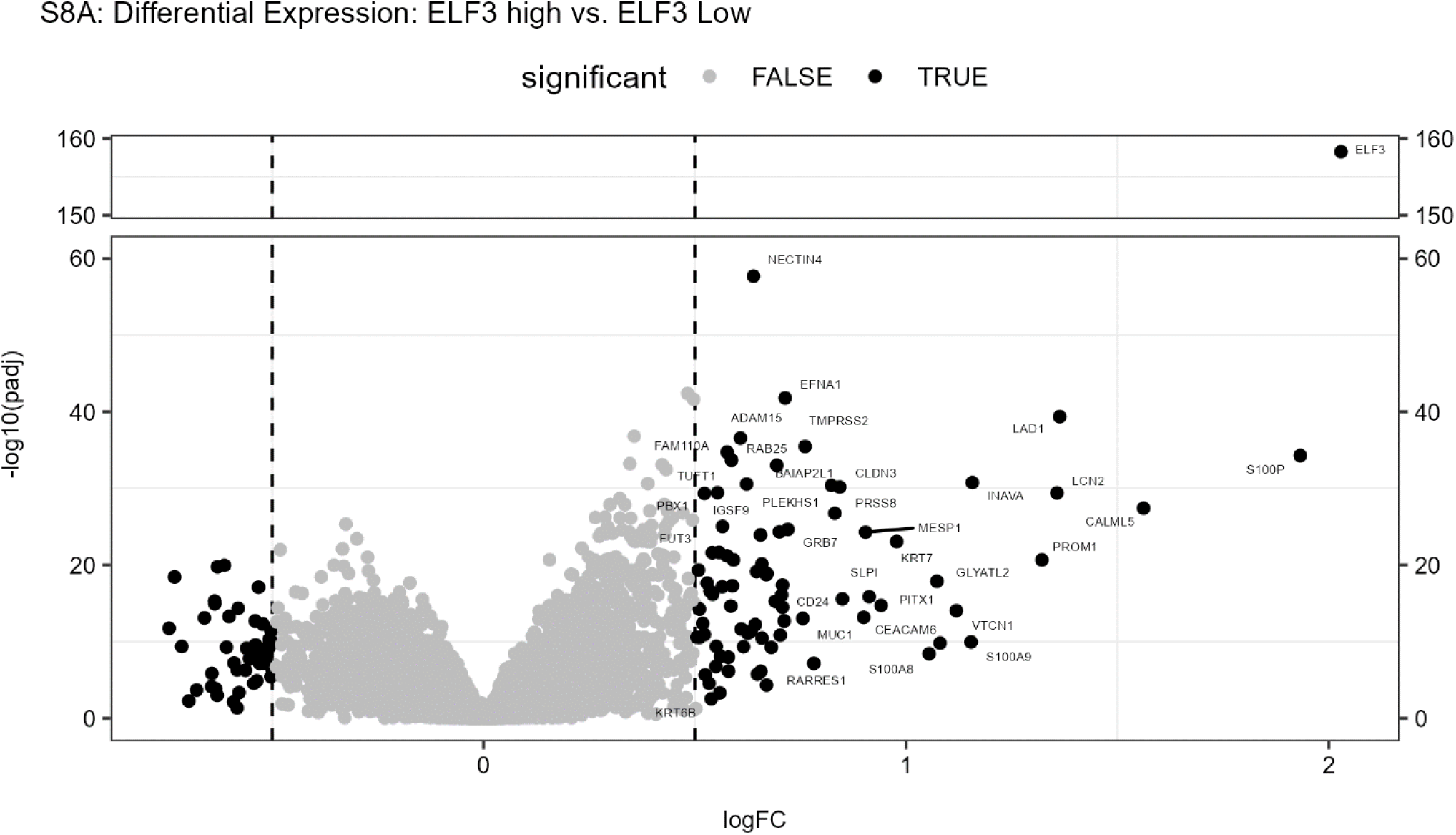
**S8A)** Volcano plot showing differentially expressed genes for the Metabric ELF3 high group versus ELF3 low group. Genes to the right (positive Log2FC) are significantly upregulated in the ELF3 high group and genes to the left (negative Log2FC) are significantly upregulated in the ELF3 low group.

## References

1. Bauer, K.R., et al., Descriptive analysis of estrogen receptor (ER)-negative, progesterone receptor (PR)-negative, and HER2-negative invasive breast cancer, the so-called triple-negative phenotype. Cancer, 2007. 109(9): p. 1721–1728.

2. Early Breast Cancer Trialists’ Collaborative, G., Comparisons between different polychemotherapy regimens for early breast cancer: meta-analyses of long-term outcome among 100 000 women in 123 randomised trials. The Lancet, 2012. 379(9814): p. 432–444.

3. Mustacchi, G. and M. De Laurentiis, *The role of taxanes in triple-negative breast cancer: literature review.* Drug Design, Development and Therapy, 2015: p. 4303.

4. Liedtke, C., et al., Response to Neoadjuvant Therapy and Long-Term Survival in Patients With Triple-Negative Breast Cancer. Journal of Clinical Oncology, 2023. 41(10): p. 1809–1815.

5. Foulkes, W.D., I.E. Smith, and J.S. Reis-Filho, Triple-Negative Breast Cancer. New England Journal of Medicine, 2010. 363(20): p. 1938–1948.

6. Schmid, P., et al., Atezolizumab and Nab-Paclitaxel in Advanced Triple-Negative Breast Cancer. New England Journal of Medicine, 2018. 379(22): p. 2108–2121.

7. Labrie, M., et al., Therapy resistance: opportunities created by adaptive responses to targeted therapies in cancer. Nature Reviews Cancer, 2022. 22(6): p. 323–339.

8. Burstein, M.D., et al., Comprehensive genomic analysis identifies novel subtypes and targets of triple-negative breast cancer. Clin Cancer Res, 2015. 21(7): p. 1688–98.

9. Spranger, S. and T.F. Gajewski, Impact of oncogenic pathways on evasion of antitumour immune responses. Nat Rev Cancer, 2018. 18(3): p. 139–147.

10. Baillie, K.E. and P.C. Stirling, Beyond Kinases: Targeting Replication Stress Proteins in Cancer Therapy. Trends Cancer, 2021. 7(5): p. 430–446.

11. Nguyen, C.D.K. and C. Yi, YAP/TAZ Signaling and Resistance to Cancer Therapy. Trends Cancer, 2019. 5(5): p. 283–296.

12. Lawson, D.A., et al., Tumour heterogeneity and metastasis at single-cell resolution. Nature Cell Biology, 2018. 20(12): p. 1349–1360.

13. Aibar, S., et al., SCENIC: single-cell regulatory network inference and clustering. Nat Methods, 2017. 14(11): p. 1083–1086.

14. Obradovic, A., et al., Single-cell protein activity analysis identifies recurrence-associated renal tumor macrophages. Cell, 2021. 184(11): p. 2988–3005 e16.

15. Ferro, A., et al., Blue intensity matters for cell cycle profiling in fluorescence DAPI-stained images. Lab Invest, 2017. 97(5): p. 615–625.

16. Zasadil, L.M., et al., Cytotoxicity of paclitaxel in breast cancer is due to chromosome missegregation on multipolar spindles. Sci Transl Med, 2014. 6(229): p. 229ra43.

17. Speyer, C.L., et al., Riluzole synergizes with paclitaxel to inhibit cell growth and induce apoptosis in triple-negative breast cancer. Breast Cancer Res Treat, 2017. 166(2): p. 407–419.

18. Lee, K.M., et al., Class III β-tubulin, a marker of resistance to paclitaxel, is overexpressed in pancreatic ductal adenocarcinoma and intraepithelial neoplasia. Histopathology, 2007. 51(4): p. 539–546.

19. Stengel, C., et al., Class III beta-tubulin expression and in vitro resistance to microtubule targeting agents. Br J Cancer, 2010. 102(2): p. 316–24.

20. Sawant, K.V., et al., Chemokine CXCL1 mediated neutrophil recruitment: Role of glycosaminoglycan interactions. Sci Rep, 2016. 6: p. 33123.

21. Cambier, S., M. Gouwy, and P. Proost, The chemokines CXCL8 and CXCL12: molecular and functional properties, role in disease and efforts towards pharmacological intervention. Cell Mol Immunol, 2023. 20(3): p. 217–251.

22. Xiong, X., et al., CXCL8 in Tumor Biology and Its Implications for Clinical Translation. Front Mol Biosci, 2022. 9: p. 723846.

23. Wang, N., et al., CXCL1 derived from tumor-associated macrophages promotes breast cancer metastasis via activating NF-kappaB/SOX4 signaling. Cell Death Dis, 2018. 9(9): p. 880.

24. Gross, S.M., et al., A multi-omic analysis of MCF10A cells provides a resource for integrative assessment of ligand-mediated molecular and phenotypic responses. Commun Biol, 2022. 5(1): p. 1066.

25. Krishna, B.M., et al., Notch signaling in breast cancer: From pathway analysis to therapy. Cancer Lett, 2019. 461: p. 123–131.

26. Arico, E., et al., Type I Interferons and Cancer: An Evolving Story Demanding Novel Clinical Applications. Cancers (Basel), 2019. 11(12).

27. Jorgovanovic, D., et al., Roles of IFN-gamma in tumor progression and regression: a review. Biomark Res, 2020. 8: p. 49.

28. Chiu, Y.H., J.B. Macmillan, and Z.J. Chen, RNA polymerase III detects cytosolic DNA and induces type I interferons through the RIG-I pathway. Cell, 2009. 138(3): p. 576–91.

29. Yu, L. and P. Liu, Cytosolic DNA sensing by cGAS: regulation, function, and human diseases. Signal Transduct Target Ther, 2021. 6(1): p. 170.

30. Keenan, A.B., et al., ChEA3: transcription factor enrichment analysis by orthogonal omics integration. Nucleic Acids Research, 2019. 47(W1): p. W212–W224.

31. Bahrami, S. and F. Drablos, Gene regulation in the immediate-early response process. Adv Biol Regul, 2016. 62: p. 37–49.

32. Jefferies, C.A., Regulating IRFs in IFN Driven Disease. Front Immunol, 2019. 10: p. 325.

33. Guler, R., et al., Batf2 differentially regulates tissue immunopathology in Type 1 and Type 2 diseases. Mucosal Immunol, 2019. 12(2): p. 390–402.

34. Liu, J., et al., Batf3(+) DCs and type I IFN are critical for the efficacy of neoadjuvant cancer immunotherapy. Oncoimmunology, 2019. 8(2): p. e1546068.

35. Rehwinkel, J. and M.U. Gack, RIG-I-like receptors: their regulation and roles in RNA sensing. Nat Rev Immunol, 2020. 20(9): p. 537–551.

36. Garces de Los Fayos Alonso, I., et al., The Role of Activator Protein-1 (AP-1) Family Members in CD30-Positive Lymphomas. Cancers (Basel), 2018. 10(4).

37. Otero, M., et al., E74-like factor 3 (ELF3) impacts on matrix metalloproteinase 13 (MMP13) transcriptional control in articular chondrocytes under proinflammatory stress. J Biol Chem, 2012. 287(5): p. 3559–72.

38. Kim, T.H., C. Kern, and H. Zhou, Knockout of IRF7 Highlights its Modulator Function of Host Response Against Avian Influenza Virus and the Involvement of MAPK and TOR Signaling Pathways in Chicken. Genes (Basel), 2020. 11(4).

39. Li, T., et al., DDIT3 and KAT2A Proteins Regulate TNFRSF10A and TNFRSF10B Expression in Endoplasmic Reticulum Stress-mediated Apoptosis in Human Lung Cancer Cells. J Biol Chem, 2015. 290(17): p. 11108–18.

40. Belanova, A.A., et al., Effects of JUN and NFE2L2 knockdown on oxidative status and NFE2L2/AP-1 targets expression in HeLa cells in basal conditions and upon sub-lethal hydrogen peroxide treatment. Mol Biol Rep, 2019. 46(1): p. 27–39.

41. Mirzaei, H., et al., The AP-1 pathway; A key regulator of cellular transformation modulated by oncogenic viruses. Rev Med Virol, 2020. 30(1): p. e2088.

42. Gross, S.M., et al., Analysis and modeling of cancer drug responses using cell cycle phase-specific rate effects. Nat Commun, 2023. 14(1): p. 3450.

43. Spencer, S.L., et al., The proliferation-quiescence decision is controlled by a bifurcation in CDK2 activity at mitotic exit. Cell, 2013. 155(2): p. 369–83.

44. Luker, K.E., et al., Overexpression of IRF9 Confers Resistance to Antimicrotubule Agents in Breast Cancer Cells1. Cancer Research, 2001. 61(17): p. 6540–6547.

45. Stringer, C., et al., Cellpose: a generalist algorithm for cellular segmentation. Nature Methods, 2021. 18(1): p. 100–106.

46. Mohammadi, F., et al., A lineage tree-based hidden Markov model quantifies cellular heterogeneity and plasticity. Commun Biol, 2022. 5(1): p. 1258.

47. Palmer, A.C. and P.K. Sorger, Combination Cancer Therapy Can Confer Benefit via Patient-to-Patient Variability without Drug Additivity or Synergy. Cell, 2017. 171(7): p. 1678–1691 e13.

48. Curtis, C., et al., The genomic and transcriptomic architecture of 2,000 breast tumours reveals novel subgroups. Nature, 2012. 486(7403): p. 346–52.

49. Symmans, W.F., et al., Long-Term Prognostic Risk After Neoadjuvant Chemotherapy Associated With Residual Cancer Burden and Breast Cancer Subtype. J Clin Oncol, 2017. 35(10): p. 1049–1060.

50. Foo, J. and F. Michor, Evolution of acquired resistance to anti-cancer therapy. J Theor Biol, 2014. 355: p. 10–20.

51. Nussinov, R., C.J. Tsai, and H. Jang, Anticancer drug resistance: An update and perspective. Drug Resist Updat, 2021. 59: p. 100796.

52. Lukow, D.A., et al., Chromosomal instability accelerates the evolution of resistance to anti-cancer therapies. Dev Cell, 2021. 56(17): p. 2427–2439 e4.

53. Abu Samaan, T.M., et al., Paclitaxel’s Mechanistic and Clinical Effects on Breast Cancer. Biomolecules, 2019. 9(12).

54. Hashemi, M., et al., EMT mechanism in breast cancer metastasis and drug resistance: Revisiting molecular interactions and biological functions. Biomed Pharmacother, 2022. 155: p. 113774.

55. Heiser, L.M., et al., Subtype and pathway specific responses to anticancer compounds in breast cancer. Proc Natl Acad Sci U S A, 2012. 109(8): p. 2724–9.

56. Barretina, J., et al., The Cancer Cell Line Encyclopedia enables predictive modelling of anticancer drug sensitivity. Nature, 2012. 483(7391): p. 603-7.

57. Yang, W., et al., Genomics of Drug Sensitivity in Cancer (GDSC): a resource for therapeutic biomarker discovery in cancer cells. Nucleic Acids Res, 2013. 41(Database issue): p. D955–61.

58. Chan, Y.T., et al., CRISPR-Cas9 library screening approach for anti-cancer drug discovery: overview and perspectives. Theranostics, 2022. 12(7): p. 3329–3344.

59. Guillon, J., et al., Chemotherapy-induced senescence, an adaptive mechanism driving resistance and tumor heterogeneity. Cell Cycle, 2019. 18(19): p. 2385–2397.

60. Kim, C.J., et al., Nuclear morphology predicts cell survival to cisplatin chemotherapy. Neoplasia, 2023. 42: p. 100906.

61. Pargett, M., et al., Single-Cell Imaging of ERK Signaling Using Fluorescent Biosensors. Methods Mol Biol, 2017. 1636: p. 35–59.

62. Netterfield, T.S., et al., Biphasic JNK-Erk signaling separates the induction and maintenance of cell senescence after DNA damage induced by topoisomerase II inhibition. Cell Syst, 2023. 14(7): p. 582–604 e10.

63. Ender, P., et al., Spatiotemporal control of ERK pulse frequency coordinates fate decisions during mammary acinar morphogenesis. Dev Cell, 2022. 57(18): p. 2153–2167 e6.

64. Grant, G.D., et al., Accurate delineation of cell cycle phase transitions in living cells with PIP-FUCCI. Cell Cycle, 2018. 17(21-22): p. 2496–2516.

65. Wang, Y., et al., Real-time imaging of senescence in tumors with DNA damage. Sci Rep, 2019. 9(1): p. 2102.

66. Balderstone, L.A., et al., Development of a fluorescence-based cellular apoptosis reporter. Methods Appl Fluoresc, 2018. 7(1): p. 015001.

67. Nam, A.S., R. Chaligne, and D.A. Landau, Integrating genetic and non-genetic determinants of cancer evolution by single-cell multi-omics. Nat Rev Genet, 2021. 22(1): p. 3–18.

68. Schiff, P.B., J. Fant, and S.B. Horwitz, Promotion of microtubule assembly in vitro by taxol. Nature, 1979. 277(5698): p. 665–667.

69. Weaver, B.A., How Taxol/paclitaxel kills cancer cells. Mol Biol Cell, 2014. 25(18): p. 2677–81.

70. Khalaf, K., et al., Aspects of the Tumor Microenvironment Involved in Immune Resistance and Drug Resistance. Front Immunol, 2021. 12: p. 656364.

71. Deepak, K.G.K., et al., Tumor microenvironment: Challenges and opportunities in targeting metastasis of triple negative breast cancer. Pharmacol Res, 2020. 153: p. 104683.

72. Ma, L., et al., Tumor Cell Biodiversity Drives Microenvironmental Reprogramming in Liver Cancer. Cancer Cell, 2019. 36(4): p. 418–430 e6.

73. Pitt, J.M., et al., Targeting the tumor microenvironment: removing obstruction to anticancer immune responses and immunotherapy. Ann Oncol, 2016. 27(8): p. 1482–92.

74. Xiao, Y. and D. Yu, Tumor microenvironment as a therapeutic target in cancer. Pharmacol Ther, 2021. 221: p. 107753.

75. Hu, J., et al., Regulation of tumor immune suppression and cancer cell survival by CXCL1/2 elevation in glioblastoma multiforme. Science Advances, 2021. 7(5): p. eabc2511.

76. Yang, M., et al., Tumour-associated neutrophils orchestrate intratumoural IL-8-driven immune evasion through Jagged2 activation in ovarian cancer. Br J Cancer, 2020. 123(9): p. 1404–1416.

77. Heidemann, J., et al., Angiogenic effects of interleukin 8 (CXCL8) in human intestinal microvascular endothelial cells are mediated by CXCR2. J Biol Chem, 2003. 278(10): p. 8508–15.

78. Choi, H.S., et al., Disruption of the NF-kappaB/IL-8 Signaling Axis by Sulconazole Inhibits Human Breast Cancer Stem Cell Formation. Cells, 2019. 8(9).

79. Rhodes, D.R., et al., Mining for regulatory programs in the cancer transcriptome. Nat Genet, 2005. 37(6): p. 579–83.

80. Margolin, A.A., et al., ARACNE: an algorithm for the reconstruction of gene regulatory networks in a mammalian cellular context. BMC Bioinformatics, 2006. 7 **Suppl 1**(Suppl 1): p. S7.

81. Wang, K., et al., Genome-wide identification of post-translational modulators of transcription factor activity in human B cells. Nat Biotechnol, 2009. 27(9): p. 829–39.

82. Subbalakshmi, A.R., et al., The ELF3 transcription factor is associated with an epithelial phenotype and represses epithelial-mesenchymal transition. J Biol Eng, 2023. 17(1): p. 17.

83. Enfield, K.S.S., et al., Epithelial tumor suppressor ELF3 is a lineage-specific amplified oncogene in lung adenocarcinoma. Nat Commun, 2019. 10(1): p. 5438.

84. Horie, M., et al., An integrative epigenomic approach identifies ELF3 as an oncogenic regulator in ASCL1-positive neuroendocrine carcinoma. Cancer Sci, 2023. 114(6): p. 2596–2608.

85. Archer, L.K., et al., ETS transcription factor ELF3 (ESE-1) is a cell cycle regulator in benign and malignant prostate. FEBS Open Bio, 2022. 12(7): p. 1365–1387.

86. Fina, E., et al., Gene signatures of circulating breast cancer cell models are a source of novel molecular determinants of metastasis and improve circulating tumor cell detection in patients. J Exp Clin Cancer Res, 2022. 41(1): p. 78.

87. Mermel, C.H., et al., GISTIC2.0 facilitates sensitive and confident localization of the targets of focal somatic copy-number alteration in human cancers. Genome Biology, 2011. 12(4): p. R41.

88. Mesquita, B., et al., Frequent copy number gains at 1q21 and 1q32 are associated with overexpression of the ETS transcription factors ETV3 and ELF3 in breast cancer irrespective of molecular subtypes. Breast Cancer Res Treat, 2013. 138(1): p. 37–45.

89. Riemenschneider, M.J., C.B. Knobbe, and G. Reifenberger, Refined mapping of 1q32 amplicons in malignant gliomas confirms MDM4 as the main amplification target. Int J Cancer, 2003. 104(6): p. 752–7.

90. Veerakumarasivam, A., et al., High-resolution array-based comparative genomic hybridization of bladder cancers identifies mouse double minute 4 (MDM4) as an amplification target exclusive of MDM2 and TP53. Clin Cancer Res, 2008. 14(9): p. 2527–34.

91. Francois, M., P. Donovan, and F. Fontaine, Modulating transcription factor activity: Interfering with protein-protein interaction networks. Semin Cell Dev Biol, 2020. 99: p. 12–19.

92. Ngamcherdtrakul, W. and W. Yantasee, siRNA therapeutics for breast cancer: recent efforts in targeting metastasis, drug resistance, and immune evasion. Transl Res, 2019. 214: p. 105–120.

93. Wolpaw, A.J., et al., Drugging the “Undruggable” MYCN Oncogenic Transcription Factor: Overcoming Previous Obstacles to Impact Childhood Cancers. Cancer Res, 2021. 81(7): p. 1627–1632.

94. Butler, A., et al., Integrating single-cell transcriptomic data across different conditions, technologies, and species. Nature Biotechnology, 2018. 36(5): p. 411–420.

95. Satija, R., et al., Spatial reconstruction of single-cell gene expression data. Nature Biotechnology, 2015. 33(5): p. 495–502.

96. Yu, G., et al., clusterProfiler: an R Package for Comparing Biological Themes Among Gene Clusters. OMICS: A Journal of Integrative Biology, 2012. 16(5): p. 284–287.

97. Dudel, C. and M. Myrskylä, Estimating the number and length of episodes in disability using a Markov chain approach. Population Health Metrics, 2020. 18(1).

98. Cerami, E., et al., The cBio cancer genomics portal: an open platform for exploring multidimensional cancer genomics data. Cancer Discov, 2012. 2(5): p. 401–4.

99. de Bruijn, I., et al., Analysis and Visualization of Longitudinal Genomic and Clinical Data from the AACR Project GENIE Biopharma Collaborative in cBioPortal. Cancer Res, 2023. 83(23): p. 3861–3867.

100. Gao, J., et al., Integrative analysis of complex cancer genomics and clinical profiles using the cBioPortal. Sci Signal, 2013. 6(269): p. pl1.

